# Delivery of oligonucleotides to bone marrow to modulate ferrochelatase splicing in a mouse model of Erythropoietic Protoporphyria

**DOI:** 10.1101/2020.02.14.949297

**Authors:** François Halloy, Pavithra S. Iyer, Paulina Ćwiek, Alice Ghidini, Jasmin Barman-Aksözen, Nicole Wildner-Verhey van Wijk, Alexandre Theocharides, Elisabeth I. Minder, Xiaoye Schneider-Yin, Daniel Schümperli, Jonathan Hall

## Abstract

Erythropoietic protoporphyria (EPP) is a rare genetic disease in which patients experience acute phototoxic reactions after sunlight exposure. It is caused by a deficiency in ferrochelatase (*FECH*) in the heme biosynthesis pathway. Most patients exhibit a loss-of-function mutation in *trans* to an allele bearing a SNP that favours aberrant splicing of transcripts. One viable strategy for EPP is to deploy splice-switching oligonucleotides (SSOs) to increase FECH synthesis, whereby an increase of a few percent would provide therapeutic benefit. However, successful application of SSOs in bone marrow cells is not described. Here, we show that SSOs comprising methoxyethyl-chemistry increase FECH levels in cells. We conjugated one SSO to three prototypical targeting groups and administered them to a mouse model of EPP in order to study their biodistribution, their metabolic stability and their FECH splice-switching ability. The SSOs exhibited distinct distribution profiles, with increased accumulation in liver, kidney, bone marrow and lung. However, they also underwent substantial metabolism, mainly at their linker groups. An SSO bearing a cholesteryl group increased levels of correctly spliced FECH transcript by 80% in the bone marrow. The results provide a promising approach to treat EPP and other disorders originating from splicing dysregulation in the bone marrow.

## INTRODUCTION

Erythropoietic protoporphyria (EPP; OMIM # 177000) is a rare autosomal recessive disorder (1), that is caused by deficiency in the heme biosynthesis enzyme ferrochelatase (*FECH*, EC <u>4.99.1.1</u>) (**Fig. 1a**) (2,3). During heme biosynthesis, FECH catalyses the incorporation of iron into protoporphyrin IX (PPIX) to form heme. With over 80 % of heme produced in red blood cells progenitors, the bone marrow is the major site of biosynthesis. Kidneys and liver are also minor sites of heme biosynthesis (4). A lack of FECH means that PPIX continuously accumulates during erythroid maturation, resulting in PPIX circulating in red blood cells of the bloodstream. Upon short exposure to visible light, PPIX produces reactive oxygen species leading to lipid peroxidation, cell membrane damage and inflammation. This causes patients to suffer from burn-like injuries of the endothelial cells of the blood vessels and the surrounding tissue, associated with acute severe pain; some will develop hepatic complications with deposits of PPIX crystals and are at risk of liver failure (5,6).

**Figure 1.**
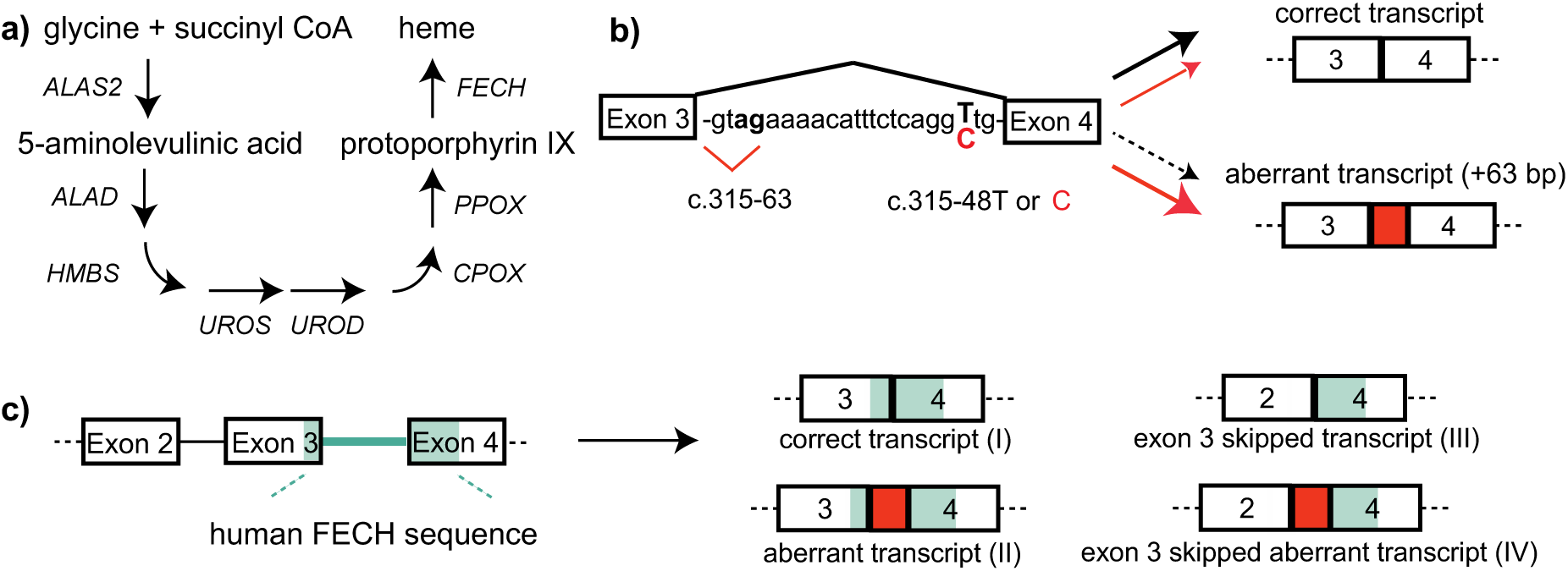
A single-nucleotide polymorphism causes aberrant splicing of *FECH* mRNA. **a)** Biosynthesis of heme from glycine and succinyl CoA comprises eight sequential steps occurring in the cytosol and in the mitochondria. FECH is the last enzyme of the pathway and catalyzes the incorporation of iron into protoporphyrin IX. EPP patients present a dramatic decrease of FECH, resulting in reduced heme production and accumulation of protoporphyrin IX in cells. **b)** FECH pre-mRNA contains a weak cryptic splice site in intron 3 (c.315-63.) close to a single nucleotide polymorphism (c.315-48T in healthy individuals). Utilization of this splice site yields a transcript which is a substrate for NMD. This aberrant splicing is increased by the c.315-48C variant which is present in trans to a hypomorphic *FECH* allele in 95% of EPP patients and in a few percent of the general population. **c)** The transgenic murine *Emi* model shows the aberrant splicing of human patients, but additionally skips exon 3 in most of the FECH transcripts.

EPP in almost all patients is associated with an alteration in both *FECH* alleles. On one, *FECH* expression is attenuated by a deleterious mutation (7); on the other, a single nucleotide polymorphism (SNP) in intron 3 (c.315-48T>C) bolsters the use of a cryptic 3’ splice site, that produces an aberrant mRNA incorporating additional sequence preceding exon 4 (**Fig. 1b**). Stop codons in this sequence result in degradation of the transcript by nonsense-mediated mRNA decay (NMD) (3). The combined effects of these alleles are reduced levels (>65%) of FECH activity, which renders patients symptomatic. As the c.315-48C allele turns a carrier of a *loss-of-function* (LoF) allele from asymptomatic to symptomatic, a splice-switching oligonucleotide (SSO) is a viable means to restore sufficient FECH expression for haplo-sufficiency in patients. SSOs composed of LNA (locked nucleic acids) and morpholino chemistries are reported to correct the aberrant splicing of the FECH mRNA *ex vivo* in patients’ blood cells (8-10), but this approach has not yet been described *in vivo*.

SSOs are oligonucleotides of about 20 nucleotides (nt) in length, that bind to a pre-mRNA and modulate its splicing (11,12). Several are in development to treat rare diseases caused by aberrant RNA maturation, reading-frame shifts, or exon skipping. The most prominent example is nusinersen, an approved treatment of spinal muscular atrophy (SMA) that mediates exon inclusion in the *SMN2* pre-mRNA (13,14). Nusinersen is composed of phosphorothioated (PS)-2’-*O-*methoxyethyl (MOE)-ribonucleotides (15), which endow the drug with high metabolic stability and extended distribution *in vivo*. PS-MOE oligonucleotides traffic to the liver, kidneys and spleen at therapeutically-useful concentrations after systemic administration (16,17). They have also been detected in bone marrow in rodents, though SSOs of various chemistries were inactive in the bone marrow of a reporter mouse model (18), possibly due to poor cellular-uptake and/or to the high turnover of red blood cell progenitors (19). This obstacle impacts an SSO strategy for EPP, but may be addressable thanks to advances in the use of conjugated groups for organ-targeted delivery (e.g. GalNAc sugars (20)).

We recently developed a transgenic mouse model for human EPP (*Emi* mouse) (21), incorporating the c.315-63 cryptic splice site of *FECH* from human intron 3. The mouse displays the aberrant splicing of exon 4; however, for reasons unknown, exon 3 is deleted from most transcripts (**Fig. 1c)**. Hence, SSOs that correct the aberrant splicing of the transgene will not increase levels of FECH activity, nor attenuate levels of circulationg PPIX. Here, we describe a 22-nt PS-MOE SSO which binds to the c.315-48 SNP and corrects splicing of *FECH* transcripts in cells and in the *Emi* mouse. Using different linker strategies, we conjugated the SSO to three targeting moieties: thiocholesterol (22,23), stearic acid (24) and a bone marrow-homing peptide (25). We first assessed their metabolic stability *in vitro*, and then their distribution, metabolism and splice-switching activity in Emi/wt heterozygous mice using mass spectrometry and chemical ligation PCR methods. Whereas the oligonucleotide moiety was metabolically stable, the conjugated groups underwent degradation *in vitro* and *in vivo*, albeit through different pathways. The conjugates also altered *in vivo*-distribution and splice-switching activity of the SSO. The parent SSO was detected in all tissues tested, and showed splice-switching of *FECH* RNA in liver, kidney and lung. The cholesterol group increased liver transport by seven-fold, but this SSO was not more potent than the parent SSO. The stearic-acid conjugate increased liver delivery by a lesser degree, but raised *FECH* transcript levels there by 60%. Notably, the homing-peptide increased bone marrow accumulation by seven-fold, but it hardly affected *FECH* splicing, whereas the cholesterol group, which did not improve transport to bone marrow, increased levels of the correct *FECH* transcript by 80%.

This study of splice-switching oligonucleotide conjugates complements recent investigations of the pharmacokinetic and pharmacodynamic properties of antisense and siRNA conjugates (26,27). The results reinforce the notion that increased delivery does not necessarily translate to improved potency. They also advance the prospect of an oligonucleotide-based therapy for correction of the *FECH* splicing defect at the origin of EPP.

## MATERIAL AND METHODS

### Small scale synthesis of MOE-PS splice-switching oligonucleotides

Small scale oligonucleotide synthesis was conducted on a MerMade 12 synthesizer (BioAutomation Inc) on a 50 nmole scale on 500 Å UnyLinker CPG (ChemGenes) with standard synthesis conditions. DMT cleavage was accomplished with a solution of 3% dichloroacetic acid in dichloromethane (V/V). 2’-*O*-MOE phosphoramidites (ThermoFisher Scientific) were prepared at 0.08 M concentration in dry acetonitrile. 5-(Benzylthio)-1H-tetrazole (Carbosynth) was used at 0.24 M concentration in dry acetonitrile to activate the phosphoramidites for coupling. Sulfurization was carried out with a 0.05 M solution of 3-((N,N-dimethylaminomethylidene)amino)-3H-1,2,4-dithiazole-5-thione (Glen Resarch #40-4037) in a 1:1 mixture of pyridine/acetonitrile. Capping of failed sequences was achieved with acetic anhydride in tetrahydrofuran (THF) and 16% N-methylimidazole in THF. Following cycle parameters were used: Deblock 2 × 60 s; Coupling 2 × 180 s; Sulfurization 1 × 150 s; Capping 2 × 50 s. Analytical data are provided in **Fig. S1**.

### Synthesis of Chemical Ligation-qPCR primers

Primers were synthesized on a MerMade 12 synthesizer following a similar procedure with minor modifications to that described in the initial publication (28). Analytical data are provided in **Fig. S2**.

### Large scale synthesis of ORN1 and of conjugates ORN_st_, ORN_84_ and ORN_ch_

Large scale synthesis was conducted on a MerMade 12 synthesizer with minor changes compared to the small-scale procedure: sulfurization reagent was prepared at 0.1 M instead of 0.05 M. Modified cycle parameters were used: Deblock 2 × 60 s; Coupling 1 × 600 s; Sulfurization 1 × 600 s; Capping 2 × 100 s (see also Supplementary Methods). Analytical data of batches used for *in vivo* work are provided in **Fig S3**.

### Oligonucleotide screening in the COS-7 FECH-C minigene system

COS-7 cells were kept in culture at 37°C, 5% CO_2_. Cells were plated at 140’000 cells / well in a 24-well plate in 1 ml DMEM-GlutaMax (ThermoFisher #10566016) containing 10 % fetal bovine serum for 18 h prior to plasmid transfection. 700 ng of the FECH-C minigene (29) was transfected using XtremeGENE HP DNA transfection reagent (Sigma-Aldrich #XTGHP-RO) as per manufacturer’s instructions with a ratio of 2.5 µl transfection reagent per µg plasmid DNA. Oligonucleotides were transfected 4 h after the plasmid using Lipofectamine™ 2000 (ThermoFisher, #11668027) following manufacturer’s instructions. 24 h prior to lysis and RNA extraction, emetine (Sigma-Aldrich #E2375) was added in each desired well at a final concentration of 3 µM.

Cells were washed with Dubelcco’s phosphate-buffered saline (DPBS) and RNA extraction was carried out using Qiagen RNEasy kit according to manufacturer’s instructions with the inclusion of two DNAse-treatment steps (Qiagen #74106, #79254). cDNA was prepared using the High Capacity cDNA RT kit (Applied Biosystems #4374966) in accordance with the manufacturer’s instructions. PCR primers specific for the minigene were used: Forward 5’-GC TCT CTA CCT GGT GTG TGG -3’, Reverse 5’-GAC AAT TCA TCC AGC AGC TTC -3’. A PCR mix containing 1 µl FW primer (10 µM stock), 1 µl RV primer (10 µM stock), 1 µl dNTPs mix (Promega #U1515), 5 µl 10X DreamTaq Buffer (ThermoFisher #EP0703), 0.25 µl DreamTaq DNA Polymerase (ThermoFisher #EP0703) and 40.75 µl ultrapure water was prepared and mixed with 1 µl of cDNA solution diluted 1/3 in water. PCR program: 95°C, 1 min; 30 cycles (95°C, 30s – 60°C, 30s – 68°C, 30s); 68°C, 5 min. PCR products were loaded on a 2% agarose gel and bands quantified using ImageJ software. Ratios were expressed as: intensity of the aberrant amplicon band / (intensities of aberrant + correct bands).

### Endogenous splicing correction and correct *FECH* transcript evaluation in K562 cells

K562 cells were kept in culture at 37°C, 5% CO_2_. Cells were plated at 200’000 cells / well in a 24-well plate in 0.5 ml IMDE (Sigma #I3390) containing 10 % fetal bovine serum and 4 mM L-glutamine (Sigma #G7513). Oligonucleotides were transfected 1.5 h after plating using Lipofectamine™ 2000 (ThermoFisher, #11668027) following manufacturer’s instructions. Emetine (Sigma-Aldrich #E2375) was added in each desired well at a final concentration of 3 µM 2.5 h hours after oligonucleotide treatment and 22 h before lysis and RNA extraction.

Cells were washed with DPBS and RNA extraction was carried out using Qiagen RNEasy kit according to manufacturer’s instructions (Qiagen #74106, #79254). cDNA was generated using PrimeScript RT kit (Takara #RR037A). For semi-quantitative RT-PCR; PCR primers specific for human *FECH* exon 2-exon 4 transcripts were used: Forward 5’-CAC AGA AAC AGC CCA GCA TG -3’, Reverse 5’-GAC AAT TCA TCC AGC AGC TTC -3’. A PCR mix containing 1.25 µl FW primer (10 µM stock), 1.25 µl RV primer (10 µM stock), 0.125 µl dNTPs mix (Promega #U1515), 0.25 µl 5X Q5 Polymerase Buffer (NEB #M0491), 0.25 µl Q5 DNA Polymerase (NEB #M0491) and 15.75 µl ultrapure water was prepared and mixed with 1 µl of cDNA solution. PCR program: 98°C, 30s; 33 cycles (98°C, 10s – 58°C, 20s – 72°C, 2s); 72°C, 2 min. PCR products were loaded on a 2% agarose gel and bands quantified using ImageJ software. Ratios were expressed as: intensity of the aberrant amplicon band / (intensities of aberrant + correct bands). Biological replicates are presented in **Fig S4**. For RT-qPCR, a primer pair for correctly spliced *FECH* transcript was used. A mix containing 4 µl SYBR Mix 2X, 0.33 µl primer pair (stock of 10 µM each in water) and 3.17 µl ultrapure water was prepared and mixed with 0.5 µl cDNA solution. Primers were purchased from Microsynth (Balgach, Switzerland) and stored as pairs at 10 µM each. Samples were run in technical triplicates and GAPDH was used as housekeeping control. Fold change calculations for the correctly spliced FECH transcript were made by comparing each treatment group with the mock group. FECH FW primer: 5’- CCT ATT CAG AAT AAG CTG GCA CC -3’; FECH RV primer: 5’- CCT GCT TGG AAG TCC ATA TCT TG -3’. GAPDH FW primer: 5’- AGG TCG GTG TGA ACG GAT TTG -3’; GAPDH RV primer: 5’- TGT AGA CCA TGT AGT TGA GGT CA -3’. Statistical analysis (multiple *t-*tests) was conducted with GraphPad Prism v7. software.

### Western Blotting experiments in K562 cells

K562 cells were grown as above. Cells were plated at 100’000 cells / well in a 24-well plate in 1 ml medium and oligonucleotides transfected 1.5 h after plating using Lipofectamine™ 2000. K562 cells were lysed 24 h or 48 h after treatment in 100 µl RIPA buffer (ThermoFisher Scientific) supplemented with protease inhibitor (cOmplete, Roche). Lysates were incubated on ice for 30 min prior to brief needle sonication (25 % amplitude, 6 s pulse), followed by centrifugation at 12’000 g for 15 min. Supernatants were collected and protein concentration was assessed with the Pierce BCA protein assay kit (ThermoFisher Scientific). 10 µg of protein from each sample was denaturated in Laemmli buffer (BioRad) containing 7% β-mercaptoethanol, and loaded onto 4-20% pre-cast TGX gels (BioRad) for electrophoresis. Protein transfer onto nitrocellulose membranes (GE Life Sciences) was carried out for 90 min at 25 W in a Mini Trans-Blot Electrophoretic Transfer Cell (BioRad). Membranes were blocked for 3 h with 5% w/v nonfat dry milk in 0.05% Tween 20 in PBS (PBS-T) and were incubated overnight at 4° C with the primary antibody. Membranes were washed three times with 0.05% PBS-T, followed by incubation with the secondary antibody for 1 hour in 5% milk in 0.05% PBS-T. After washing with 0.05% PBS-T, signal was detected with the ECL Prime reagent (GE Life Sciences) in a ChemiDoc XRS+ station (BioRad).

Chemiluminescence was quantified using the ImageJ software. Ratios were calculated and graphical output and statistical analysis (one-way ANOVA) was conducted with GraphPad Prism v7. software. Biological replicates are presented in **Fig S5**. List of primary antibodies used: FECH (mouse, SantaCruz, sc-377377, 1/500 dilution), GADPH (mouse, Proteintech, 60004-1-Ig, 1/10’000 dilution) and the corresponding secondary antibody: anti-mouse (goat, Seracare, 5220-0341, 1/10’000 dilution).

### Stability assay in mouse plasma

5 nmoles of *in-vivo* ready oligonucleotides (UV purity >99%, sodium salt) were spiked in 200 µl of mouse K2-EDTA plasma (BioIVT #MSE02PLK2Y2N) and 20 µl aliquots were taken immediately after mixing, and after 5 min, 15 min, 30 min, 1 h, 2 h, 4 h, 6.5 h, 22 h and 26 h incubation and gentle shaking at 37°C. Aliquots were immediately quenched with 350 µl OTX Clarity Lysis buffer (Phenomenex #AL0-8579) and frozen at −20°C. Clean-up was conducted all at once using the Clarity® OTX Extraction kit (Phenomenex #KS0-9253) according to manufacturer’s procedure for biological fluids with the following adjustments: formulation of all buffers with ammonium acetate in substitution of sodium phosphate; dissolution of the final eluate after evaporation to dryness in 100 µl of TE buffer pH 8.0. Reconstituted samples were analysed qualitatively by LC-MS (Agilent 1200/6130 system) equipped with a Waters Acquity OST C-18 column.

### Animal injections

The oligonucleotides and oligonucleotide conjugates were injected subcutaneously into mice of the B6.CgJ-*Fech*^*tm1*.*1(FECH*)Emi*^ line (*Emi* for short) of *Emi/wt* genotype at a dose of 50 mg/kg. Four doses were administered over a period of two weeks. Weight of the mice was determined prior to each injection to monitor any adverse weight loss due to oligonucleotide injection. 24 h after the last injection, the mice were weighed and euthanized by carbon dioxide inhalation. Blood was withdrawn from the posterior vena cava of each mouse and used to prepare heparinized plasma. Liver, spleen, kidneys, lungs, heart, brain, quadriceps muscle were collected and snap frozen in liquid nitrogen. Bone marrow was isolated from femurs and tibiae by flushing the bones with cell culture medium (RPMI +10% FBS). Bone marrow cell pellets were frozen at −80°C till used for uptake studies or RNA isolation. Tissue samples were ground into a fine powder in liquid nitrogen and separate samples were immediately aliquoted for dissolution in OTX Lysis buffer (Phenomenex #AL0-8579) for uptake and metabolite study, or in Trizol (ThermoFisher #1559606) for RNA extraction. All animal experiments were conducted in accordance with local laws under the license number ZH115/16.

### CL-qPCR in tissues

The ground tissue powder was suspended in 10 volumes/weight OTX Lysis Buffer, briefly needle-sonicated and centrifuged for 30 s at 14’000 rpm. Supernatants were collected, diluted at 1/750 in ultrapure water, and used for analysis. CL-qPCR was carried out following the protocol from Boos et al.(28), with minor modifications. For each tissue and each compound, a calibration curve was generated and run along with the samples. Mix composition for Chemical Ligation: 0.1 µl PS primer (10 µM stock), 0.1 µl BPS primer (10 µM primer), 1 µl Poly A (GE Healthcare #27411001, 1 mg/ml), 1 µl 10X Buffer with MgCl_2_ (Roche #12032902001) and 5.8 µl ultrapure water, mixed with 2 µl diluted lysate. Chemical ligation was run for 1 h at 33°C. qPCR was run on a LC480 device (Roche). A qPCR mix containing 0.15 µl FW primer (10 µM stock), 0.15 µl RV primer (10 µM stock), 0.13 µl BHQ primer (28.28 µM stock), 0.15 µl dNTPs mix (ThermoFisher #10297018), 1 µl 10X Buffer with MgCl_2_ (Roche #12032902001), 0.1 µl 10X FastTaq Polymerase (Roche #12032902001) and 6.32 µl ultrapure water was prepared and mixed with 2 µl of chemical ligation reaction mixture. qPCR program was as follows: 10 min, 95°C, 50 cycles (3 s/95°C – 30 s/55°C – 10 s/72°C). Concentrations in tissues were obtained through interpolation from the calibration curves and expressed in ng oligonucleotide / mg tissue for each mouse.

### Oligonucleotide extraction from tissue lysates for LC-MS analysis

Ground tissues were suspended in 10 volumes/weight OTX Lysis Buffer, briefly needle-sonicated and centrifuged for 30 s at 14’000 rpm. Supernatants were collected for oligonucleotide clean-up and directly processed with the Clarity® OTX Extraction kit (Phenomenex #KS0-9253) according to manufacturer’s procedure for tissues with the following adjustments: formulation of all buffers with ammonium acetate in substitution of sodium phosphate; dissolution of the final eluate after evaporation to dryness in 100 µl of TE buffer pH 8.0 instead of water. Proteinase K pre-treatment was necessary for successful recovery of the oligonucleotide with liver samples coming from mice treated with cholesteryl conjugate **ORN1**_**ch**_; this pre-treatment was used for all subsequent **ORN1**_**ch**_ tissues.

### RNA extraction from mouse tissues and RT-qPCR for correct FECH transcript

The ground tissue powder for each sample was dissolved in Trizol reagent (10 to 50 mg tissue/ml Trizol) and RNA was extracted following manufacturer’s instructions. RNA samples were subjected to an extra clean-up step and DNase treatment using Qiagen RNeasy kit as per manufacturer’s instructions (Qiagen #74106, #79254). cDNA was generated using PrimeScript RT kit (Takara #RR037A). qPCR was run on a LC480 device (Roche) with the KAPA SYBR Fast qPCR kit (#KK4618). A qPCR mix containing 4 µl SYBR Mix 2X, 0.33 µl primer pair (stock of 10 µM each in water) and 3.17 µl ultrapure water was prepared and mixed with 0.5 µl cDNA solution. Primers were purchased from Microsynth (Balgach, Switzerland) and stored as pairs at 10 µM each. Samples were run in technical triplicates and GAPDH was used as housekeeping control. Fold change calculations for the FECH correctly spliced transcript were made by comparing each treatment group with the saline group. FECH FW primer: 5’- CCTATTCAGAATAAGCTGGCACC-3’; FECH RV primer: 5’- GGG GAT CCG CCT CCA ATC-3’. GAPDH FW primer: 5’-AGG TCG GTG TGA ACG GAT TTG -3’; GAPDH RV primer: 5’- TGT AGA CCA TGT AGT TGA GGT CA-3’. Statistical analysis (multiple *t-*tests) and outlier removal (ROUT method) were conducted with GraphPad Prism v7. software.

## RESULTS AND DISCUSSION

### Identification of a lead SSO for FECH RNA splicing correction

As of yet, it is difficult to predict potent SSO sequences using computational methods (30,31). Therefore, the search for an effective SSO typically begins by “walking” empirically a set of overlapping sequences across exonic and intronic regions in the region of the pathogenic mutation. This approach identified nusinersen, which binds the ISS-N1 site on SMN2 (14), and more recently, an effective SSO as a potential treatment for familial dysautonomia (32).

We scanned regions on the FECH pre-mRNA close to the c.315-48 polymorphism (**Fig 2a**; sequences in Supplementary Information) for splice-switching activity by using a set of sequence-overlapping 19-nt MOE-PS oligonucleotides. We elected to use MOE-PS SSOs because of their successful use in clinical programs (www.ionispharma.com) (33), their excellent safety record (33,34), their efficient distribution *in vivo* (16), the ease with which they can be elaborated with targeting groups or stereopure linkages (29). We employed a mini-gene assay transiently expressing the FECH-C variant in COS-7 cells, which we have described previously (21). For each SSO, we calculated the fraction of “correct” (desired) transcripts on agarose gels after work-up using RT-PCR. Since the aberrant transcript, at least for the complete FECH gene, is degraded through non-sense mediated mRNA decay (NMD) (3), we pre-treated cells with emetine, a global inhibitor of translation that prevents NMD (35) to ensure reliable measurements of aberrant transcript levels. Most SSOs showed some effect on target splicing, changing the fraction of correct transcripts by a few percent in the desired direction (**Fig 2b**). Several SSOs binding between residues 40-66 produced almost full splice correction, and activity dropped away with sequences binding upstream of position −50. In a round of optimization for increased potency, we identified two 22-nt- and one 25-nt SSOs displaying equal or better activity than the aformentioned LNA SSO (9), which we used as a control (**Fig 2c, 2d)**. From these, **ORN1** was selected as the lead SSO.

**Figure 2.**
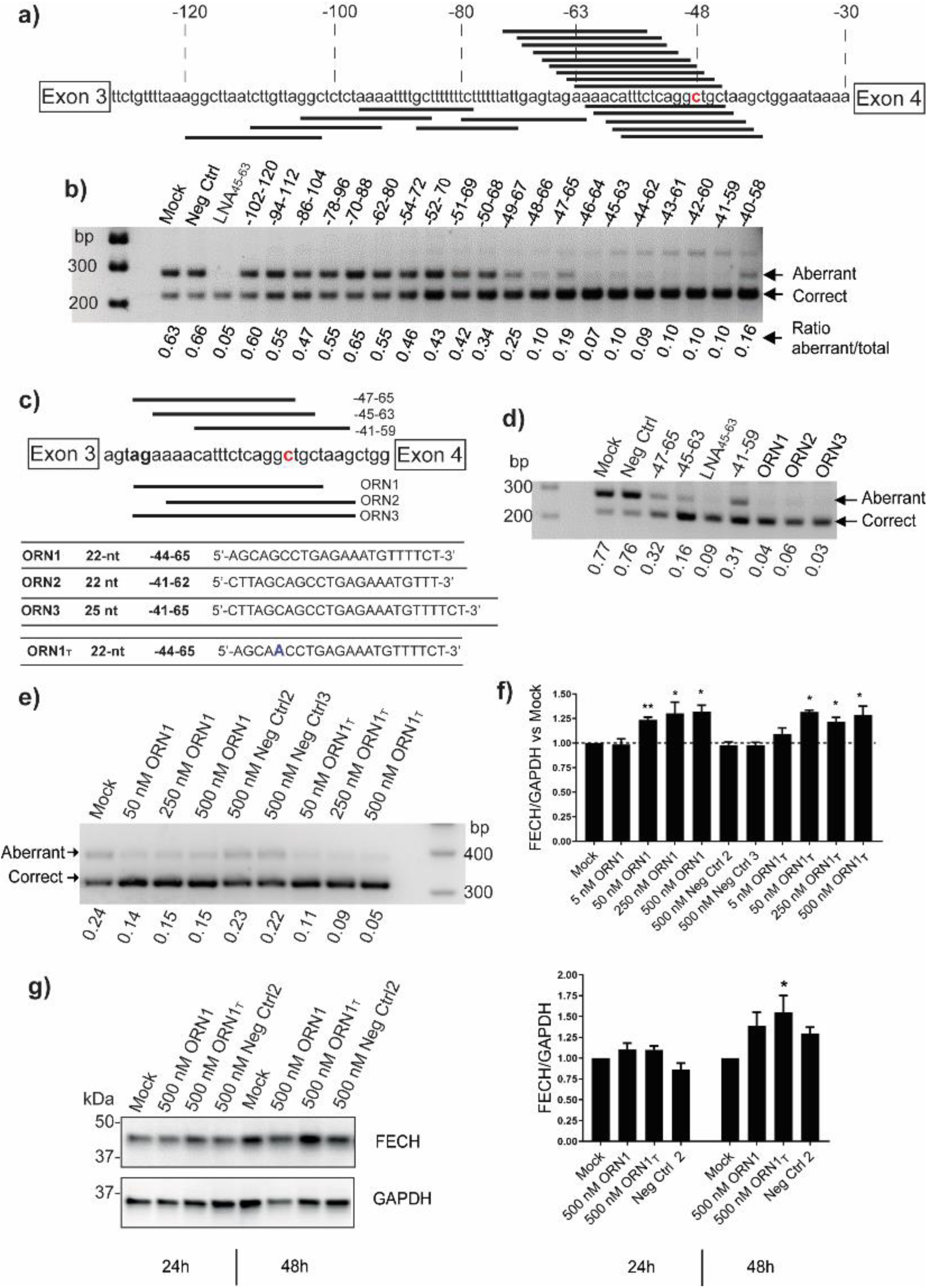
Identification of splice-switching oligonucleotides (SSOs) targeting intron 3 of FECH pre-mRNA. **a)** Binding sites for twenty 19-nt MOE-PS SSOs in the region of the *FECH* c.315-48 polymorphism and the c.315-63 aberrant splice site. **b)** Agarose gel showing relative amounts of aberrant and correct transcripts after treatment with 50 nM concentrations of SSOs in COS-7 cells expressing a FECH-C minigene; LNA_45-63_ (8) is a positive control. **c)** Lead optimization: binding sites for 22 and 25-nt SSO analogs of selected SSOs from b). **d)** Activity of **ORNs 1-3** showing complete splice correction at 10 nM concentration in the same assay. **e)** Activity of **ORN1, ORN1**_**T**_ and two randomized versions of **ORN1** in K562 cells 24 h after transfection (**ORN1** and **ORN1**_**T**_ bind FECH mRNA in K562 cells with a G-U wobble and with perfect complementarity, respectively). Aberrant transcripts were stabilized with 3 µM emetine treatment. **f) ORN1** and **ORN1**_**T**_ selectively increase levels of endogenous correctly spliced *FECH* transcripts in K562 cells as assayed by SYBR Green qPCR. n = 3 biological replicates; statistical analysis (t tests) was conducted with GraphPad Prism software: (*) = p < 0.05; (**) = p < 0.01. **g)** Representative Western Blot of FECH and GAPDH after a single transfection at 500 nM in K562 cells after 24 h or 48 h treatments; corresponding graph plot from n = 3 biological replicates; statistical analysis (one-way Anova) was conducted with GraphPad Prism software: (*) = p < 0.05 compared to mock treatments.

### Binding the c.315-48C SNP but not the aberrant splice site is sufficient for splicing correction

We noted from the initial screen that the most active SSOs covered the −48 SNP, but not necessarily the −63 aberrant splice site. We tested a library of shorter (10-nt) MOE-PS oligonucleotides (**Table S1**, Supplementary information) binding to the −63 and the −48 sites of FECH separately, in order to pinpoint the nucleotides important for the activity of the SSOs in the region of the SNP. One sequence binding at positions −45-54 altered the course of splicing significantly, and three-nucleotide shifts upstream or downstream resulted in loss of activity (**Fig. S6**). Co-treatments of cells with SSOs blocking simultaneously the splice site (−57-66) and the SNP (−45-54) did not produce synergistic effects.

We and others (3) have observed that ratios of aberrant:correct FECH transcripts vary according to the genotype, the cell type and iron availability (36). With emetine treatment to inhibit NMD of mRNA in lymphoblasts immortalized from donors, we found approximately 30% and 60% of the FECH transcripts were aberrantly spliced for the T/T (c.315-48T)- and C/C (c.315-48C)-homozygotes, respectively (not shown). We observed the same ratios in COS-7 cells transfected with the minigenes (**Fig. 2b**) and approximately 15% aberrant splicing in the K562 cell line (T/T variant). This implies that the outcome of FECH splicing may depend upon a finely-tuned balance of positive and negative splicing regulatory elements. According to Human Splicing Finder (37), the −48 SNP is embedded in predicted binding sites for hnRNPA1, SF2 and SC35 alternative splice factors, and the T>C polymorphism may create an Exon Inclusion Element (EIE) (38). An oligonucleotide binding to the −45-54 locus might lessen this activation, in a similar fashion to that described for the *cblE* type of homocystinuria, where the c.309+469T>C mutation creates an EIE and promotes the inclusion of a pseudo-exon (39).

It has not yet been possible to test **ORN1** for its effects on native FECH mRNA and protein, since transfectable patient-derived (C/C)-cells expressing low levels of FECH protein are unavailable, and we have so far been unable to generate a model cell line using CRISPR-CAS based methods. Therefore, we assayed **ORN1** in K562 erythroleukemia cells, expressing the T-variant of FECH, in which **ORN1** binds its target with a G-U wobble at the site of the polymorphism. We included a fully complementary version of the SSO (**ORN1**_**T**_) in the assay. Both SSOs showed splice-switching activity, increasing the levels of the correctly spliced transcript by 25-30% (**Fig 2f**) and FECH protein by ∼50% (**Fig. 2g**).

### Synthesis of MOE-PS SSOs conjugated with groups for targeted delivery

MOE-PS oligonucleotides accumulate in bone marrow after systemic administration in rodents (16,18) and in man (see refs in (17)), however, MOE-PS SSOs were inactive when tested in a mouse model for splicing of a reporter gene (18). Therefore, we conjugated **ORN1** with three prototypical functionalities that are reported to enhance delivery to selected tissues *in vivo* (**Fig 3a**): stearic acid which increases circulation times due to albumin binding (24) (**ORN**_**st**_); a heptapeptide residue that homes to bone marrow and binds primitive stem cells (25) (**ORN**_**84**_); and cholesterol which delivers single-stranded and double-stranded oligonucleotides to bone marrow (22,40,41) (**ORN**_**ch**_). Stearic acid was conjugated as an amide whereas the peptide and the cholesterol groups were conjugated *via* thiol-maleimide chemistry (see Supplementary Methods) (42,43).

**Figure 3.**
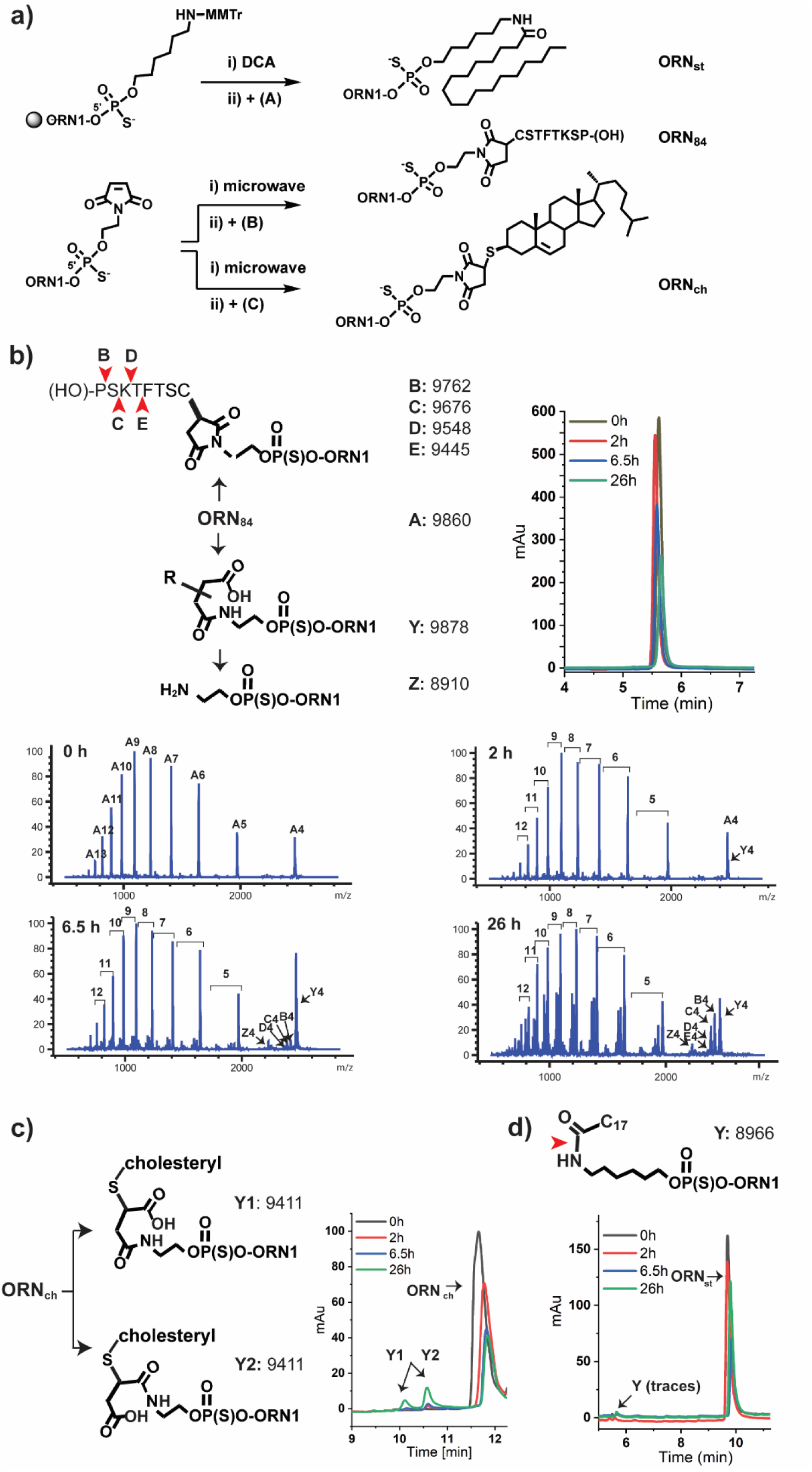
Synthesis of SSO conjugates and their metabolism in mouse plasma. **a)** Synthesis routes of MOE-PS SSO conjugates. The stearic acid conjugate **ORN**_**st**_ was obtained in two steps through coupling of a 5’-amino C_6_-phosphoramidite linker, followed by reaction with stearic N-hydroxy succinimide ester (**A**); the peptide conjugate **ORN**_**84**_ and the cholesterol conjugate (**ORN**_**ch**_) were generated in two steps using a 5’-caged maleimide modifier phosphoramidite and reaction with the cysteine-modified CD84-peptide (**B**) or thiocholesterol (**C**), respectively. (DCA: dichloroacetic acid; MMTr: monomethoxy-trityl). **b)-d) ORN**_**84**_, **ORN**_**ch**_ and **ORN**_**st**_ were incubated in mouse plasma for 26 h at 37°C. Aliquots of plasma were removed periodically, cleaned and analysed by LC-MS. **ORN1**_**84**_ underwent hydrolysis at the peptide and the succinimide linker (b). **ORN1**_**ch**_ underwent hydrolysis-mediated ring opening (c). **ORN1**_**st**_ was barely affected under the conditions (d). Red arrows indicate inferred sites of cleavage, based upon the masses of fragments in the LC-MS chromatagrams.

### *In vitro* characterization of conjugate stability in plasma and on albumin binding

MOE-PS oligonucleotides are highly resistant to nuclease-mediated degradation *in vivo* (44,45), but less is known about the metabolic stability of the conjugates and their linkers. The thiosuccinimide linker of **ORN**_**84**_ and **ORN**_**ch**_ has been widely used in the field of antibody drug conjugates but may undergo thiol-exchange reactions with cysteines of serum albumin (46) or glutathione (47). The peptide of **ORN**_**84**_ is not modified to convey stability against hydrolases. Therefore, we examined the stability of the SSOs in a plasma stability assay that mimics conditions in the bloodstream (48,49). We incubated the SSOs in mouse plasma and removed aliquots periodically for analysis by liquid-chromatography mass spectrometry (LC-MS).

**ORN1** was highly high stable in plasma, though trace amounts of a 3’ (n-1) metabolite was detected after 26 h (not shown). In contrast, all of the conjugates suffered some degradation. The peptide moiety of **ORN1**_**84**_ underwent sequential proteolysis from the C-terminus: metabolites missing one (**B**) and two amino-acids (**C**) were seen after 6.5 h, and (**D**) and (**E**) were identified after 26 h (**Fig. 3b**). The linker also underwent ring opening of the succinimide (metabolite **Y**) after 1 h, and cleavage of the amide was apparent after 6.5 h (metabolite **Z**). **ORN**_**ch**_ underwent a similar albeit slower hydrolysis of the linker, producing two products with the same mass (9411: **ORN**_**ch**_+18), which we assigned as ring-opened regioisomers (**Y1** and **Y2**) (**Fig. 3c**). The stearic acid conjugate **ORN**_**st**_ showed only trace amounts of degradation after 26 h (**Fig. 3d**).

### Distribution and metabolism of FECH SSOs in mouse tissues and organs

Recently, we engineered a murine model of EPP (*Emi*), in which intron 3 of the mouse gene was replaced by the corresponding human sequence bearing the c.315-48C polymorphism (21) (**Fig. 1c**). Homozygous *Emi/Emi* mice were not viable. However, viable *Emi/wt* mice contain one wild-type allele and one allele of the EPP genotype which produces the same aberrant/correct splicing products as the human c.315-48C allele. Unfortunately, this splicing phenotype was accompanied by an additional unanticipated splicing defect - an almost complete skipping of exon 3, and thus loss of FECH protein. Hence, the *Emi* model is unsuitable for studying effects of SSOs on the disease phenotype, but is amenable to investigations of the pharmacokinetics properties of SSOs and their effects on the amount of *FECH* transcripts containing the extra 63 nucleotide sequence (**Fig. 1c**).

SSOs were administered to *Emi*/*wt* mice twice a week over two weeks similar to other published protocols using MOE-PS oligonucleotides (16,51-53). Mice were sacrificed 24 h after the last injection. No signs of toxicity were apparent from measurements of total body weight, liver/body weight and spleen/body weight ratios as well as assessment of the general condition (not shown). Accumulation of SSOs in selected tissues was quantified using chemical ligation – qPCR (CL-qPCR) (28,50,54) (**Fig. 4b**). This technique is particularly useful for quantification of SSOs that are heavily modified and are not amplifiable by polymerases. Standard calibration curves for the four SSOs were generated in diluted lysate **(Fig S7)** and used to determine SSO concentrations in the tissues of treated mice.

**Figure 4.**
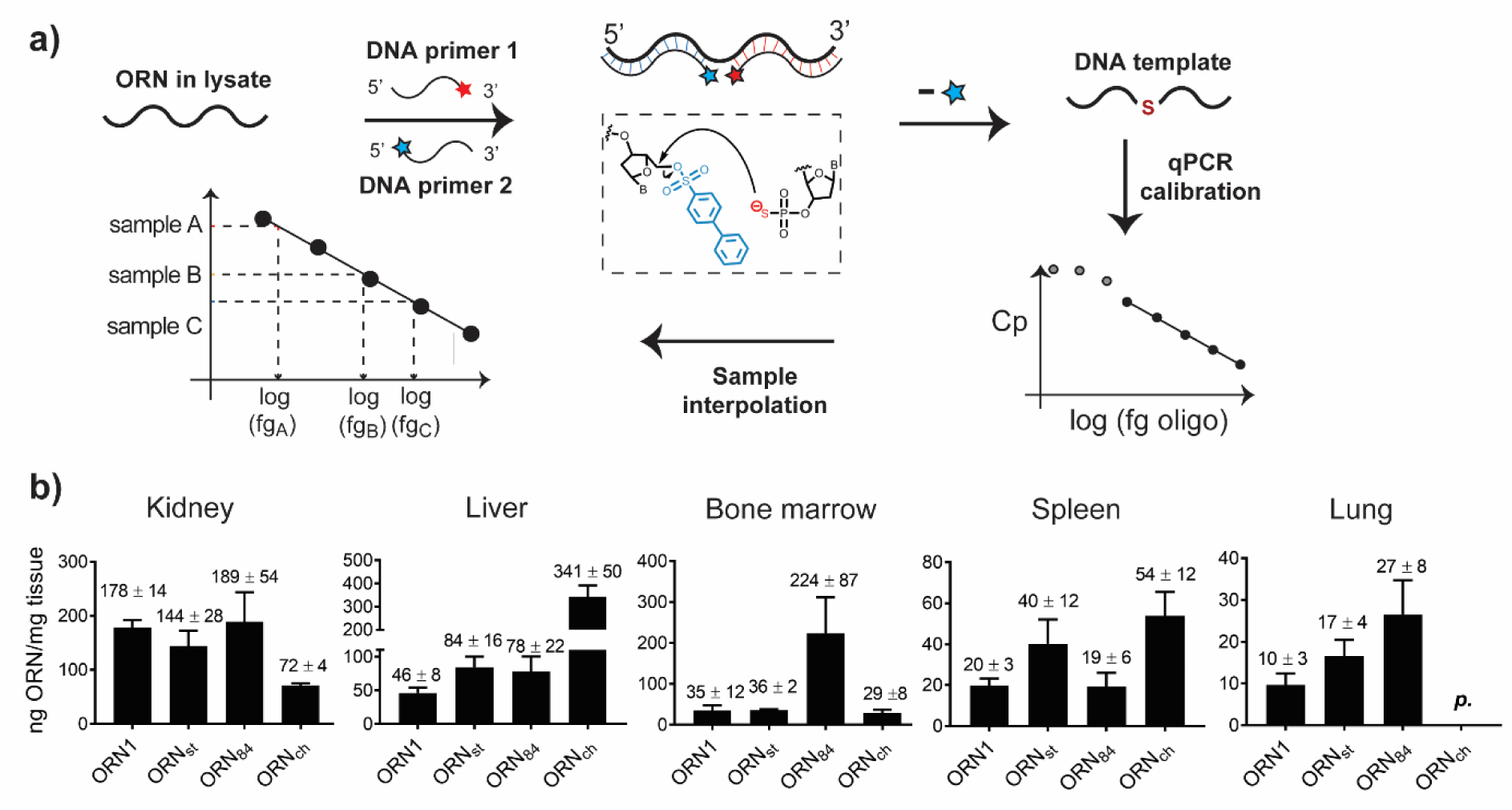
Characterization of SSOs and their accumulation in selected tissues. **a)** Principle of CL-qPCR (50). Two chemically modified PCR primers “PS” and “PBS” are added to a sample containing the chemically modified SSO and incubated for 1 h at 30°C. Hybridization of the primers to the SSO lead to formation of a DNA strand, which serves as template for quantitative PCR. **b)** Quantification using CL-qPCR of target oligonucleotides **ORN1, ORN**_**ch**_, **ORN**_**st**_ and **ORN**_**84**_ in kidney, liver, bone marrow, spleen and lung. Animals received 4 injections at 50 mg/kg in 0.9% saline over two weeks and were sacrificed 24 h after the last dose. n = 4; data expressed as mean value ± s.e.m.; p = present (detectably higher than NTC, but not quantifiable).

**ORN1** was detected in all tissues except for brain and muscle. Consistent with previous studies (55), the highest levels were found in kidney (178±14 ng/mg tissue) and liver (46±8 ng/mg tissue), followed by bone marrow (31±17 ng/mg tissue), spleen (20±3 ng/mg tissue) and lung (10±3 ng/mg tissue). Furthermore, the absolute concentrations of **ORN1** measured using CL-qPCR are in accordance with those determined using radio-labelled oligonucleotides (16). **ORN**_**st**_ showed a two-fold higher uptake in liver (84±16 ng/mg tissue), in spleen (40±12 ng/mg tissue) and lung (17±4 ng/mg tissue), and a slightly reduced accumulation in kidney. These findings are again consistent with reports that siRNAs conjugated to hydrophobic fatty acids are transported to the liver at the expense of kidney (26). **ORN**_**ch**_ showed seven-fold higher levels in liver (341±50 ng/mg tissue), a two-fold increase in spleen (54±12 ng/mg tissue) and a 60% decrease in kidney compared to **ORN1**; the presence of **ORN**_**ch**_ in lung tissue was below accurate detection. Again, this is consistent with the findings of other groups using cholesteryl-siRNAs (24,56) and single-stranded antisense-cholesteryl conjugates (57). **ORN**_**84**_ showed an approximate two-fold increase in liver (78±22 ng/mg tissue) and lung (27±8 vs. 10±3 ng/mg tissue), and markedly improved delivery to bone marrow (224±87 vs. 31±17 ng/mg tissue) in line with its previously described properties (25).

We investigated the metabolic fate of the four SSOs after repeated administration *in vivo*. Tissue samples were processed using an OTX Clarity Kit and eluates were collected from kidney, liver, spleen, lung, brain and bone marrow. The extraction protocol worked poorly for samples containing **ORN**_**ch**_, which was remedied by pre-treatment with proteinase K. Reference LC-MS spectra were generated by spiking SSOs into eluates collected from tissues from saline-treated mice. This permitted unambiguous identification and qualitative analysis of oligonucleotide-derived peaks in the chromatograms from samples from treated animals (**Fig. 5a**). Deconvolution of the mass spectra returned molecular weights from which the structures of metabolites were inferred (**Fig. 5b**; **Figs. S8-S11**).

**Figure 5.**
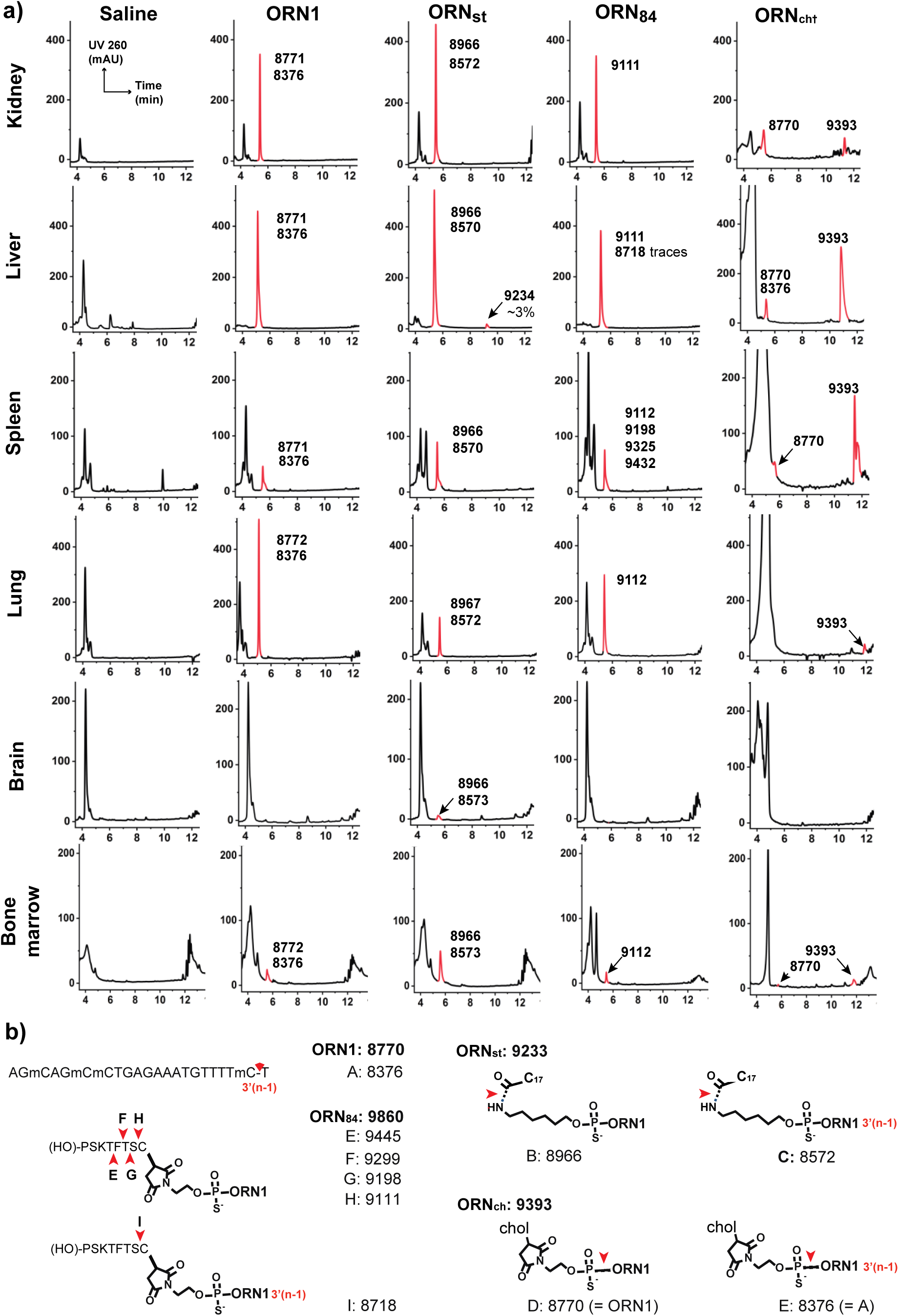
Metabolites of SSOs ORN1, ORN_st_, ORN_84_ and ORN_ch_ from tissues after administration to *Emi*/*wt* mice. **a)** Animals received 4 injections at 50 mg/kg in 0.9% saline over two weeks and were sacrificed 24 h after the last dose. Oligonucleotides were extracted from the tissue matrix using the Phenomenex Clarity OTX kit. Tissues from mice treated with **ORN1**_**ch**_ were subjected to a proteinase K digestion before processing. Detected oligonucleotide species are marked in red, next to their associated masses determined by LC-MS (see also **Figs. S8**-**S11**) b) Calculated molecular weights of parent SSOs and their inferred metabolites from LC-MS data shown in a) (C17 is the stearoyl chain; chol is the thiocholesteryl group).

Intact **ORN1** was detected in all the tissues except the brain, accompanied by 5-10% of the (n-1)-metabolite corresponding to loss of the 3’-terminal nucleotide (**A**). In contrast, all of the SSO conjugates showed fragmentation. Despite its high stability in mouse plasma, fully intact **ORN**_**st**_ was only detected in trace amounts in the liver sample: hydrolysis of the stearoyl group was seen in all tissues (**B**), together with minor amounts of the 3’(n-1)-metabolite (**C**). Since the stearoyl group of this SSO was stable in plasma (**Fig. 3d**), it seems possible that its metabolism occurred at the site of accumulation, or intracellularly. Similar to **ORN**_**st**_, **ORN**_**84**_ was not found intact in any tissues and the majority of its metabolites derived from progressive cleavage of the peptide. The major metabolite in liver, kidney, lung and bone marrow was the SSO attached to the terminal cysteine residue (**H** and **I**), whereas partial cleavage of the peptide occurred in spleen (metabolite **G**, possibly with **E** and **F**). **ORN**_**ch**_ was the most resistant of the series, and delivered a strong signal for intact **ORN**_**ch**_ from kidney, liver and spleen samples; the SSO was also detectable in lung and in bone marrow in lower amounts. Interestingly, cleavage of the PS-linkage to yield **ORN1** and its 3’-truncated fragment (**A**) was observed in samples from kidney, liver and spleen as well as bone marrow.

### Effects of SSOs on FECH RNA splicing in mouse tissues and organs

As discussed above, splice-switching activity of the SSOs in treated *Emi*/*wt* mice could not be assayed at the protein level, as the predominant transcript III (**Fig. 1c**) from the humanized allele lacks exon 3. Since the human aberrant transcript is a substrate for NMD (3), we doubted that the aberrantly spliced transcript II from the transgene could be accurately measured. Therefore, we assayed SSO activity by the increase in the correctly-spliced transcript I by using SBYR Green RT-qPCR. Splice correction of the *FECH* transgene by SSOs was observed in kidney, liver, bone marrow, spleen and lung, with varying efficiency; no splicing modulation was detected in spleen, heart and gastrocnemius muscle (**Fig. 6**). Kidney is the main site of accumulation for single-stranded PS oligonucleotides (**Fig. 4b**) (16,58).

**Figure 6.**
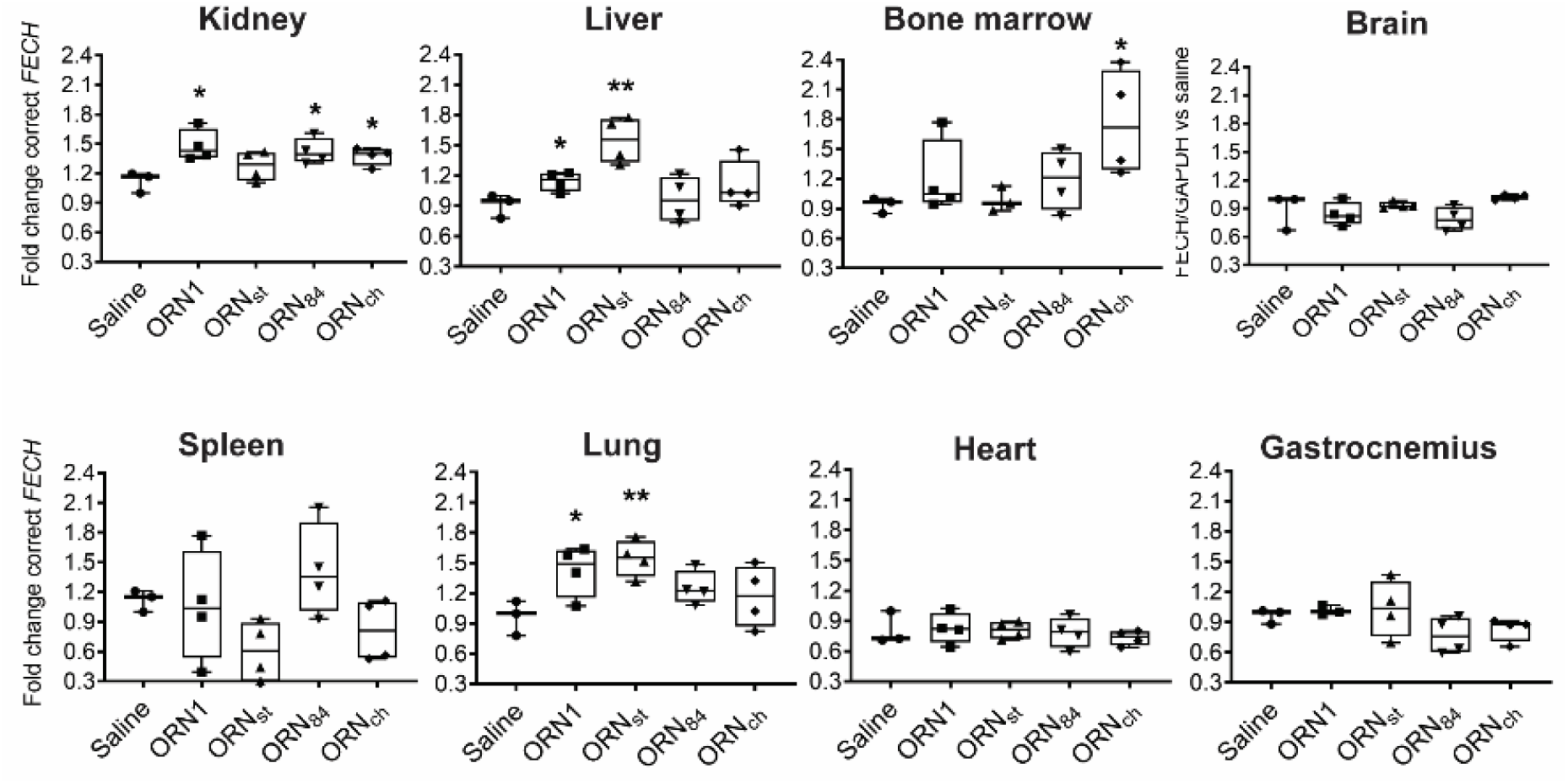
Treatment of *Emi/wt* mice with SSOs modulates splicing in selected tissues. Animals received 4 injections at 50 mg/kg in 0.9% saline over two weeks and were sacrificed 24 h after the last injection. Levels of correct FECH transcript were assessed by SYBR green RT-qPCR and normalized to GAPDH and to the saline group. n = 3 animals for the saline group; n = 4 animals for oligonucleotide treated groups; statistical analysis (t tests) was conducted with GraphPad Prism software: (*) = p < 0.05; (**) = p < 0.01 (whiskers indicate lowest and highest values; middle line represents the median).

The stearoyl conjugate **ORN**_**st**_ was most effective in liver (1.6-fold) and in lung, two tissues in which the conjugated group increased accumulation (**Fig. 4b**). **ORN**_**84**_, which was found at 6.5-fold higher levels in bone marrow (**Fig. 4b**), did not show significantly better splice-modulating activity than **ORN1** (**Fig. 6**).

From the three SSO conjugates, **ORN**_**ch**_ provided the most promising results. Surprisingly, it did not exhibit superior activity to that of **ORN1** in liver nor in kidney. This was unexpected since cholesterol conjugates are a well-known means to boost the activities of antimiR oligonucleotides in liver (22) (**ORN**_**ch**_ exhibited a 7-fold increase in liver; **Fig. 4b**). Most importantly, however, **ORN**_**ch**_ showed a 1.8-fold increase in the *FECH* transcript in bone marrow compared to **ORN1**, with a similar level of accumulation. Since most of the *FECH* transcripts in the bone marrow likely originate from erythroblasts, the increase in correctly spliced FECH mRNA strongly suggests a successful uptake of the SSO by erythroblasts.

## DISCUSSION

Loss-of-function (LoF) mutations that result in reduced or abolished protein function represent a common cause of genetic disorders. Many of these result from defective splicing events (59). It is difficult to discover or design conventional small-molecule drugs to correct defective splicing in a sequence-selective manner. However, oligonucleotides are well suited to modulate mRNA splicing since they bind with high selectivity directly to the disease-causing element. Consequently, there is considerable excitement for a wave of SSOs that are currently in pre-clinical and clinical development (32,60-62).

Splice-switching mechanisms differ fundamentally from terminating mechanisms for oligonucleotide therapies. For the latter, the drug is typically optimized to provide a maximum inhibition of the target; for the former, the objective is to increase expression of the target to counter a loss of function. For many LoF diseases, even a small increase in levels of the target protein can be therapeutically beneficial (63,64). This applies to EPP where it is thought that raising FECH protein/enzyme activity by 10-20% may be sufficient to provide therapeutic benefit to patients (8). The development of an oligonucleotide therapy for EPP is especially desireable since nearly all patients exhibit the same intronic SNP variant. On the other hand, an oligonucleotide therapy for EPP requires the delivery of the SSO into erythrocyte precursors in the bone marrow and with an efficiency to counter the rapid pace of erythrocyte biogenesis.

In this study, we identified PS-MOE SSOs that bind to the c.315-48 SNP and redirect the splicing of FECH, almost completely to the correct form in cell models. It is likely that the SSO blocks an EIE at the site of the SNP. We conjugated the SSO to three classes of targeting groups (cholesterol, fatty acids, peptides) and administered them to a mouse model of EPP in order to study their distribution, their metabolic stability and their FECH splice-switching ability. The conjugated groups exhibited distinct distribution profiles, notably increasing SSO delivery to liver, kidney, bone marrow and lung. A variety of sensitive techniques have emerged recently for the detection and quantification of oligonucleotides in biological samples (28,65,66). We employed CL-qPCR to quantify the SSOs in tissues and recorded similar values to those reported using other methods (10’s-100’s ng/mg tissue). Whereas the PS-MOE oligonucleotide was highly stable, the conjugates were heavily metabolized *in vivo*, particularly the amide linkage of **ORN**_**st**_, which was unexpected given its common use in drug conjugates (67-69). This metabolism likely occurred after clearance from circulation, since amide cleavage was not observed in the mouse plasma assay. The most robust SSO *in vivo* was **ORN**_**ch**_, which did not boost the accumulation in bone marrow but nevertheless increased levels of correctly spliced FECH transcript by 80%. It is presently unknown whether a 1.8-fold increase in FECH transcript levels would produce a therapeutically-meaningful outcome, i.e. would lower circulating PPIX. However, the experiments performed in the K562 cells are encouraging, where a 30% increase of correctly spliced RNA raised levels of FECH protein by 50% (**Fig. 2f-g**). Clarity on this question will be available from an improved mouse model of EPP (ongoing), i.e. expressing the same FECH transcripts (I and II in **Fig. 1c**) present in EPP patients.

The recent approval of several breakthrough oligonucleotide therapeutics signifies that RNA can today be considered as a mainstream class of drugs and drug targets. The oligonucleotides are single- or double-stranded, of various chemistries and they elicit their action upon well-validated mRNA targets expressed in the liver and the central nervous system. Extending the use of SSO therapies to the bone marrow, for example for EPP or other disorders, would significantly advance the oligonucleotide field.

## ACKNOWLEDGEMENTS

We thank Annamari Alitalo from the ETH Phenomics Center for useful discussions.

