## Supplementary Information for "Delivery of oligonucleotides to bone marrow to modulate ferrochelatase splicing in a mouse model of Erythropoietic Protoporphyria"

#### Supplementary figures

##### Figure S1. Oligonucleotide analytics for *in vitro* work in COS-7 and K562 cells

Samples were analysed by LC-MS (Agilent 1200/6130 system) equipped with a Waters Acquity OST C-18 column with a gradient of MeOH in 0.4 mM HFIP, 15 mM trimethylamine; flow rate: 0.3 ml/min. The target refers to the position bound on *FECH* pre-mRNA.

| Name | Seq. 5’-3’ | Target | Chemistry | MW found (g/mol) | MW calc. (g/mol) | ∆m (%) | UV purity (%) |
|---|---|---|---|---|---|---|---|
| **1** | g$^{m}$c$^{m}$ctaa$^{m}$caagattaag$^{m}$c$^{m}$ct | -102-120 | MOE-PS | 7547.22 | 7547 | 0.01 | 98 |

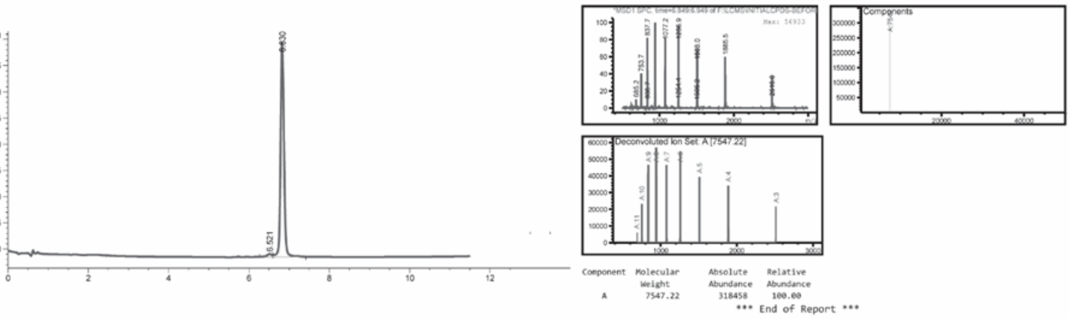

| Name | Seq. 5’-3’ | Target | Chemistry | MW found (g/mol) | MW calc. (g/mol) | ∆m (%) | UV purity (%) |
|---|---|---|---|---|---|---|---|
| **2** | tttagagag$^{m}$c$^{m}$ctaa$^{m}$caaga | -94-112 | MOE-PS | 7582.98 | 7582 | 0.01 | >99 |

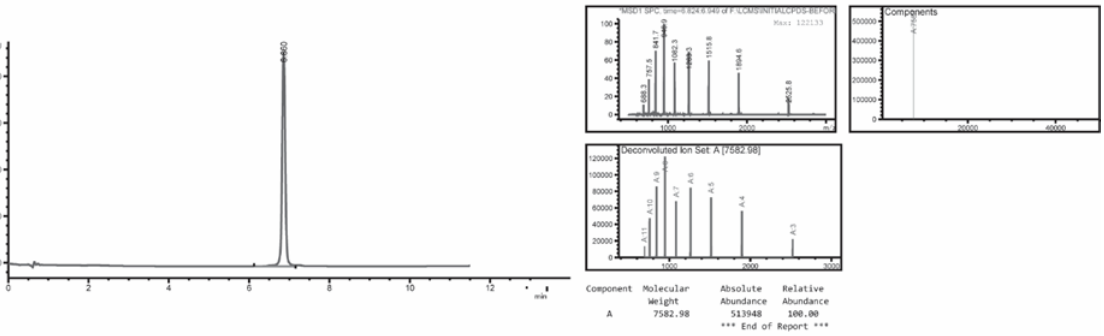

| Name | Seq. 5’-3’ | Target | Chemistry | MW found (g/mol) | MW calc. (g/mol) | ∆m (%) | UV purity (%) |
|---|---|---|---|---|---|---|---|
| **3** | ag$^{m}$caaaattttagagag$^{m}$c$^{m}$c | -86-104 | MOE-PS | 7582.45 | 7582 | 0.01 | >99 |

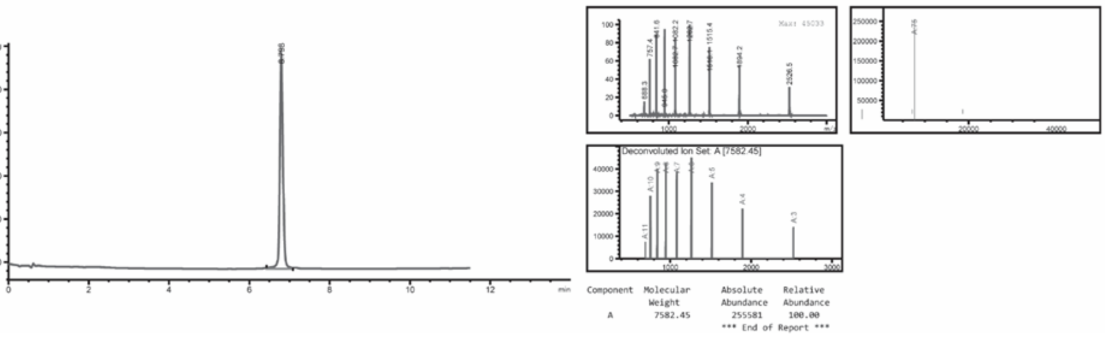

| Name | Seq. 5'-3' | Target | Chemistry | MW found (g/mol) | MW calc. (g/mol) | Δm (%) | UV purity (%) |
|---|---|---|---|---|---|---|---|
| **4** | agaaaaaaag$^{m}$caaaatttt | -78-96 | MOE-PS | 7573.13 | 7571 | 0.02 | 98 |

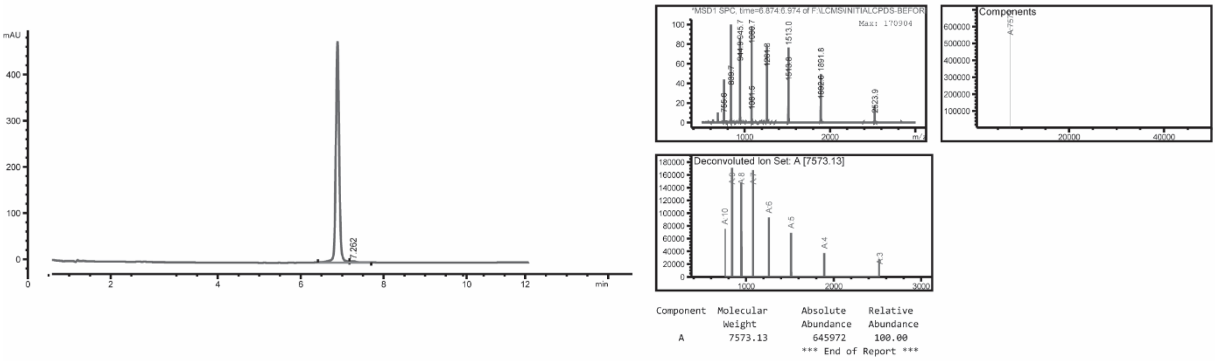

| Name | Seq. 5'-3' | Target | Chemistry | MW found (g/mol) | MW calc. (g/mol) | Δm (%) | UV purity (%) |
|---|---|---|---|---|---|---|---|
| **5** | aataaaaaagaaaaaaag$^{m}$c | -70-88 | MOE-PS | 7598.62 | 7597 | 0.02 | >99 |

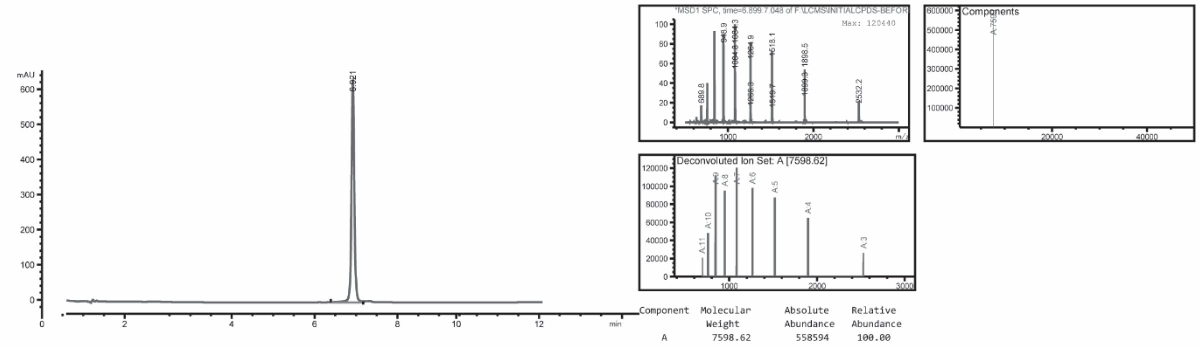

| Name | Seq. 5'-3' | Target | Chemistry | MW found (g/mol) | MW calc. (g/mol) | Δm (%) | UV purity (%) |
|---|---|---|---|---|---|---|---|
| **6** | tt$^{m}$cta$^{m}$ctcaauaaaaaaga | -62-80 | MOE-PS | 7525.52 | 7525 | 0.01 | >99 |

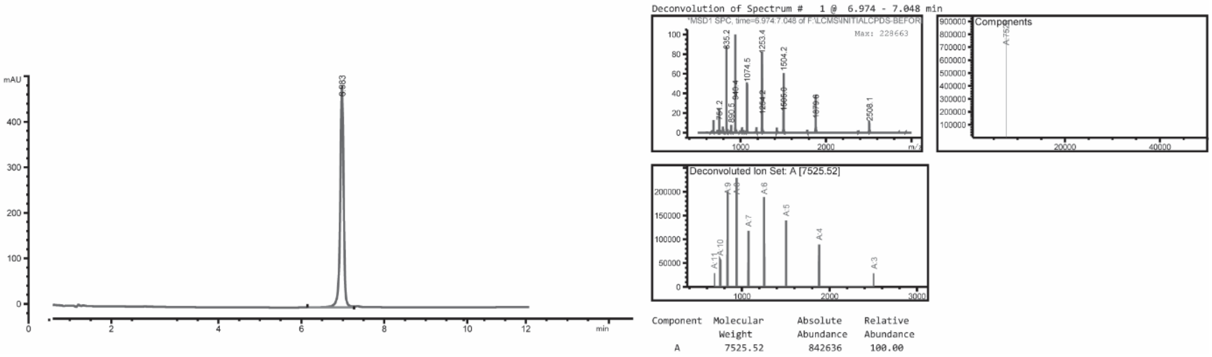

| Name | Seq. 5'-3' | Target | Chemistry | MW found (g/mol) | MW calc. (g/mol) | Δm (%) | UV purity (%) |
|---|---|---|---|---|---|---|---|
| **7** | Gaaatgttttctactcaat | -54-72 | MOE-PS | 7515.80 | 7514 | 0.01 | >99 |

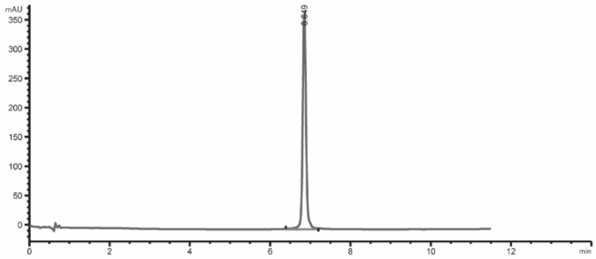
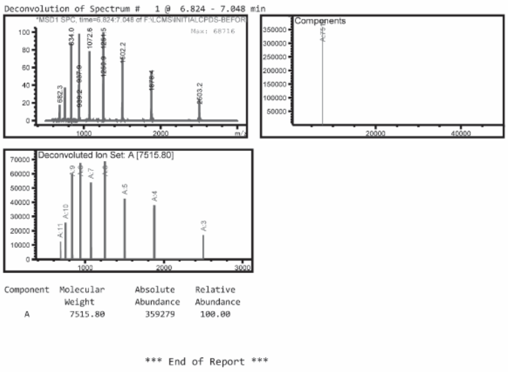

| Name | Seq. 5'-3' | Target | Chemistry | MW found (g/mol) | MW calc. (g/mol) | Δm (%) | UV purity (%) |
|---|---|---|---|---|---|---|---|
| **8** | gagaaatgtttt$^{m}$cta$^{m}$ct$^{m}$ca | -52-70 | MOE-PS | 7555.33 | 7553 | 0.01 | >99 |

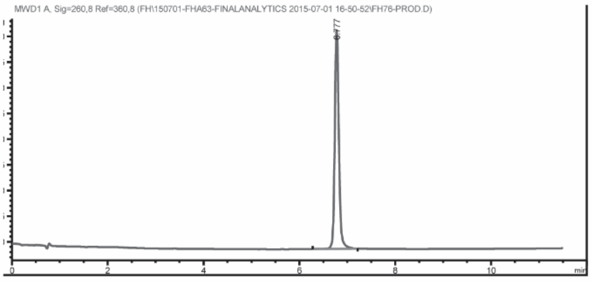

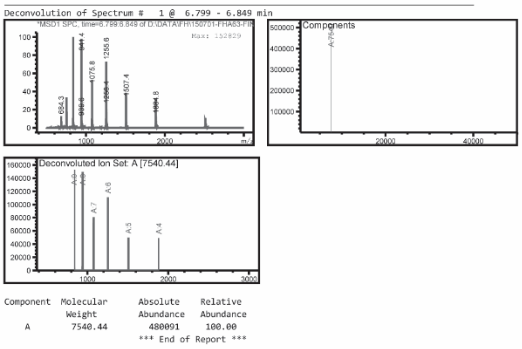

| Name | Seq. 5'-3' | Target | Chemistry | MW found (g/mol) | MW calc. (g/mol) | Δm (%) | UV purity (%) |
|---|---|---|---|---|---|---|---|
| **9** | tgagaaatgtttt$^{m}$cta$^{m}$ct$^{m}$c | -51-69 | Full MOE-PS | 7530.69 | 7529.8 | 0.01 | >99 |

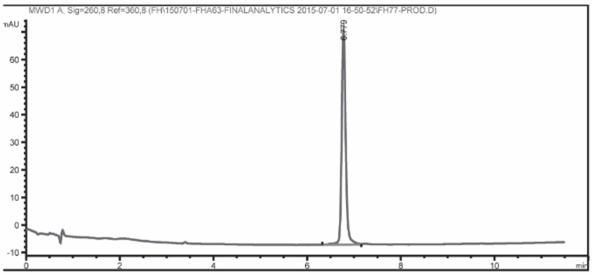

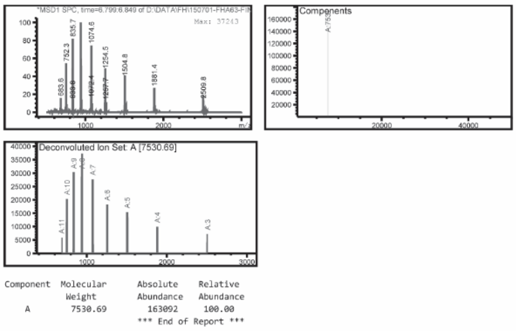

| Name | Seq. 5’-3’ | Target | Chemistry | MW found (g/mol) | MW calc. (g/mol) | Δm (%) | UV purity (%) |
|---|---|---|---|---|---|---|---|
| **10** | $^{m}$ctgagaaatgtttt$^{m}$cta$^{m}$ct | -50-68 | Full MOE-PS | 7530.69 | 7529.8 | 0.01 | >99 |

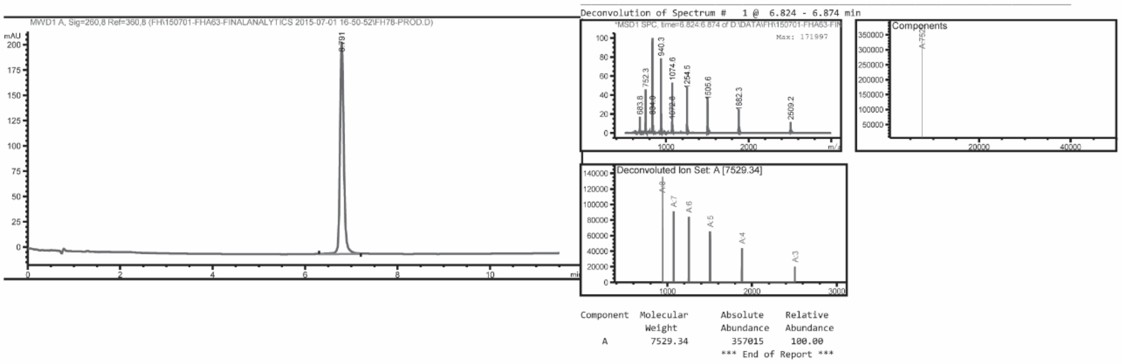

| Name | Seq. 5’-3’ | Target | Chemistry | MW found (g/mol) | MW calc. (g/mol) | Δm (%) | UV purity (%) |
|---|---|---|---|---|---|---|---|
| **11** | $^{m}$c$^{m}$ctgagaaatgtttt$^{m}$cta$^{m}$c | -49-67 | Full MOE-PS | 7531.24 | 7528.8 | 0.03 | >99 |

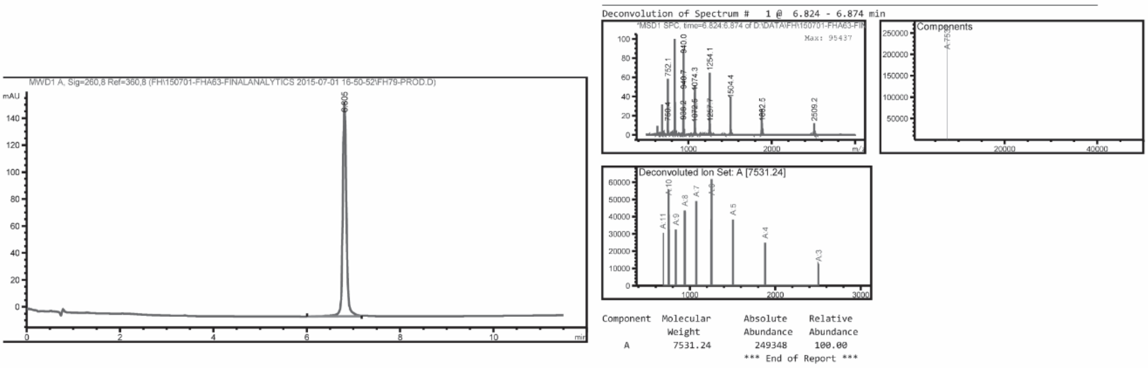

| Name | Seq. 5’-3’ | Target | Chemistry | MW found (g/mol) | MW calc. (g/mol) | Δm (%) | UV purity (%) |
|---|---|---|---|---|---|---|---|
| **12** | g$^{m}$c$^{m}$ctgagaaatgtttt$^{m}$cta | -48-66 | Full MOE-PS | 7555.40 | 7554.8 | 0.01 | >99 |

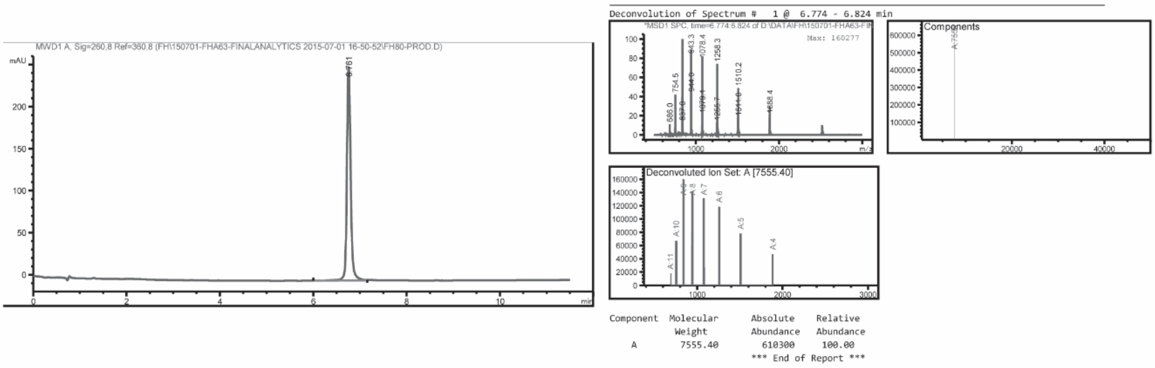

| Name | Seq. 5'-3' | Target | Chemistry | MW found (g/mol) | MW calc. (g/mol) | Δm (%) | UV purity (%) |
|---|---|---|---|---|---|---|---|
| **13** | ag$^{m}$c$^{m}$ctgagaaatgtttt$^{m}$ct | -47-65 | MOE-PS | 7556.59 | 7555 | 0.01 | >99 |

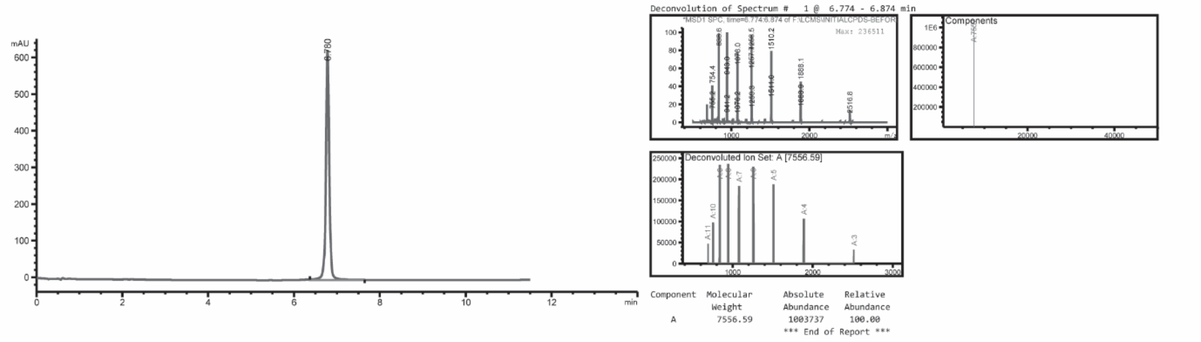

| Name | Seq. 5'-3' | Target | Chemistry | MW found (g/mol) | MW calc. (g/mol) | Δm (%) | UV purity (%) |
|---|---|---|---|---|---|---|---|
| **14** | $^{m}$cag$^{m}$c$^{m}$ctgagaaatgtttt$^{m}$c | -46-64 | MOE-PS | 7555.33 | 7553 | 0.01 | >99 |

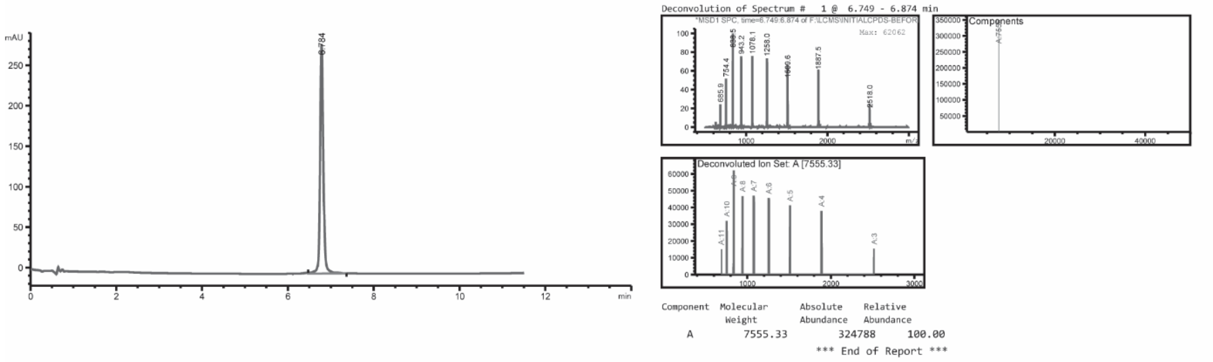

| Name | Seq. 5'-3' | Target | Chemistry | MW found (g/mol) | MW calc. (g/mol) | Δm (%) | UV purity (%) |
|---|---|---|---|---|---|---|---|
| **15** | g$^{m}$cag$^{m}$c$^{m}$ctgagaaatgtttt | -45-63 | MOE-PS | 7580.48 | 7580 | 0.01 | >99 |

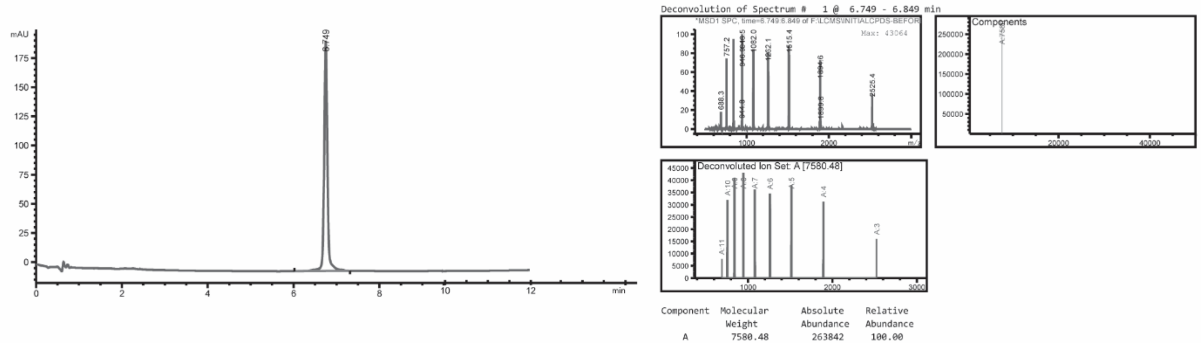

| Name | Seq. 5'-3' | Target | Chemistry | MW found (g/mol) | MW calc. (g/mol) | Δm (%) | UV purity (%) |
|---|---|---|---|---|---|---|---|
| **16** | ag$^{m}$cag$^{m}$c$^{m}$ctgagaaatgttt | -44-62 | Full MOE-PS | 7588.02 | 7588.9 | 0.01 | >99 |

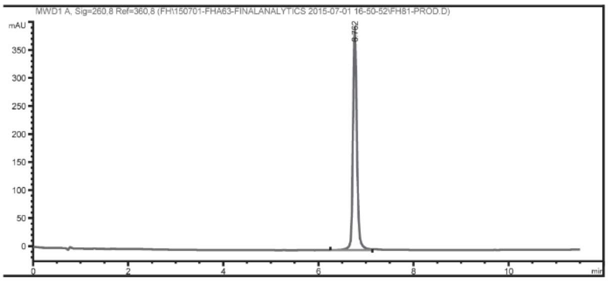

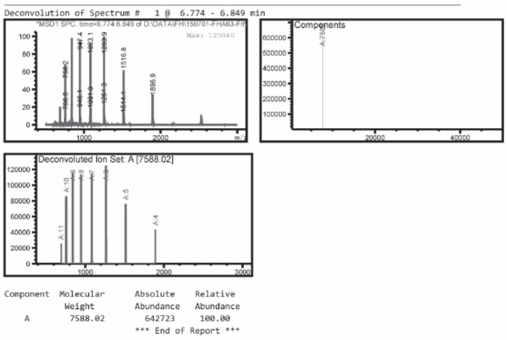

| Name | Seq. 5'-3' | Target | Chemistry | MW found (g/mol) | MW calc. (g/mol) | Δm (%) | UV purity (%) |
|---|---|---|---|---|---|---|---|
| **17** | tag$^{m}$cag$^{m}$c$^{m}$ctgagaaatgtt | -43-61 | Full MOE-PS | 7589.52 | 7588.9 | 0.01 | >99 |

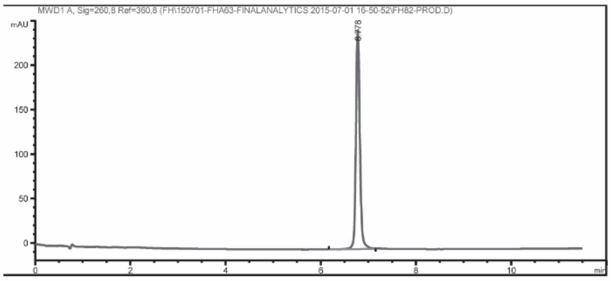

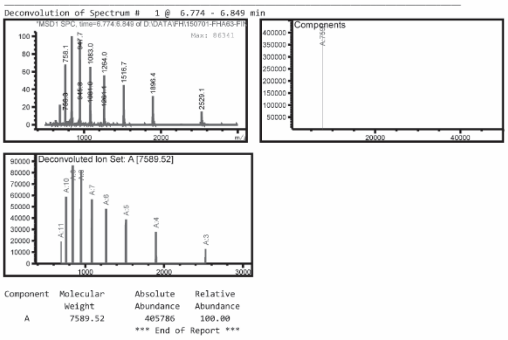

| Name | Seq. 5'-3' | Target | Chemistry | MW found (g/mol) | MW calc. (g/mol) | Δm (%) | UV purity (%) |
|---|---|---|---|---|---|---|---|
| **18** | ttag$^{m}$cag$^{m}$c$^{m}$ctgagaaatgt | -42-60 | Full MOE-PS | 7589.09 | 7588.9 | <0.01 | >99 |

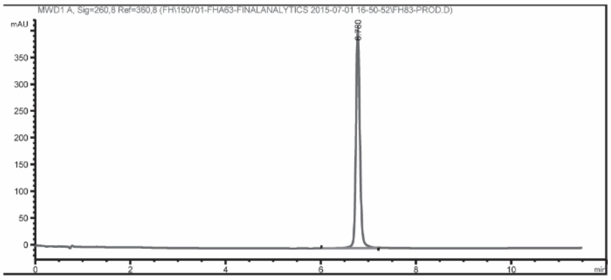

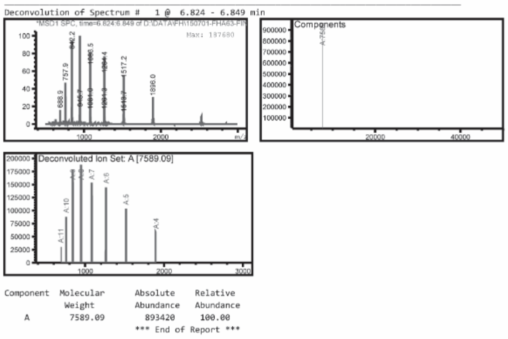

| Name | Seq. 5'-3' | Target | Chemistry | MW found (g/mol) | MW calc. (g/mol) | Δm (%) | UV purity (%) |
|---|---|---|---|---|---|---|---|
| **19** | $^{m}$cttag$^{m}$cag$^{m}$c$^{m}$ctgagaaatg | -41-59 | Full MOE-PS | 7588.54 | 7587.9 | 0.01 | 98.7 |

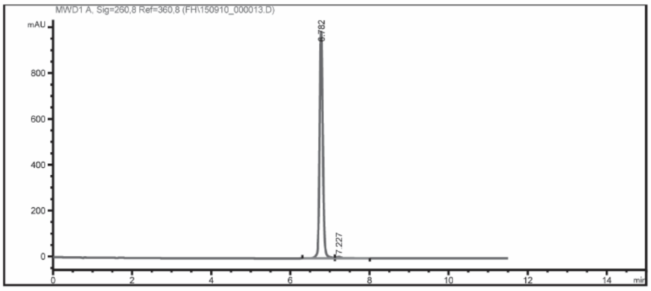

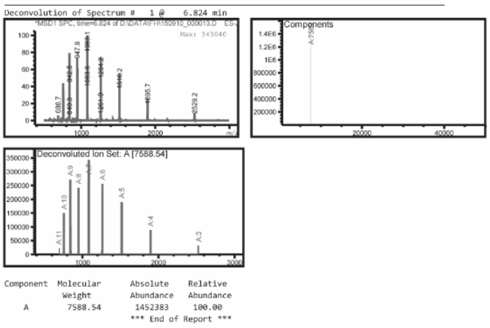

| Name | Seq. 5'-3' | Target | Chemistry | MW found (g/mol) | MW calc. (g/mol) | Δm (%) | UV purity (%) |
|---|---|---|---|---|---|---|---|
| **20** | g$^{m}$cttag$^{m}$cag$^{m}$c$^{m}$ctgagaaat | -40-58 | Full MOE-PS | 7588.83 | 7587.9 | 0.01 | >99 |

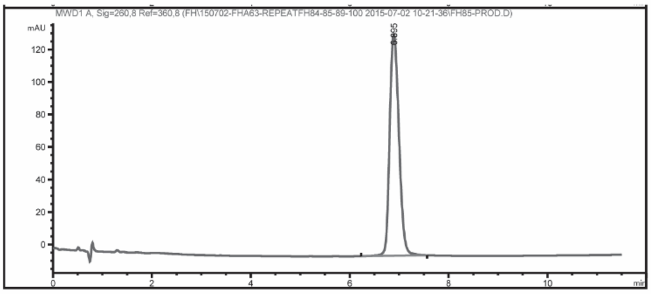

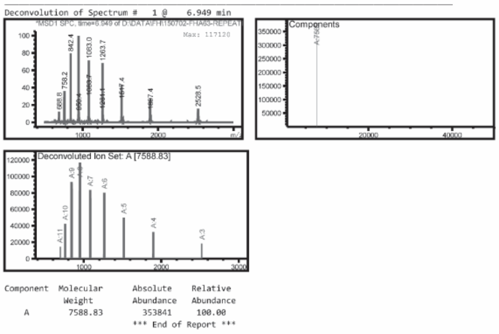

| | Name | Seq. 5'-3' | Target | Chemistry | MW found (g/mol) | MW calc. (g/mol) | Δm (%) | UV purity (%) |
|---|---|---|---|---|---|---|---|---|
| **Neg Ctrl 1** | **21** | gttattg$^{m}$ctat$^{m}$cgaag$^{m}$cag | none | Full MOE-PS | 7583.00 | 7579.9 | 0.04 | >99 |

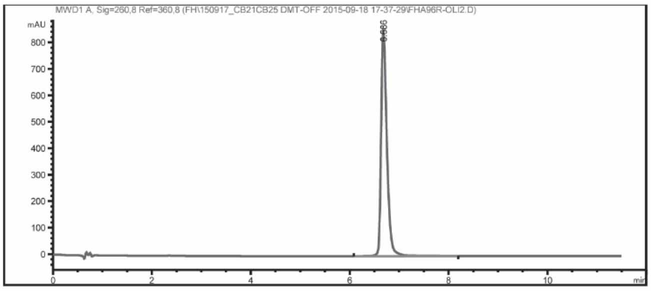

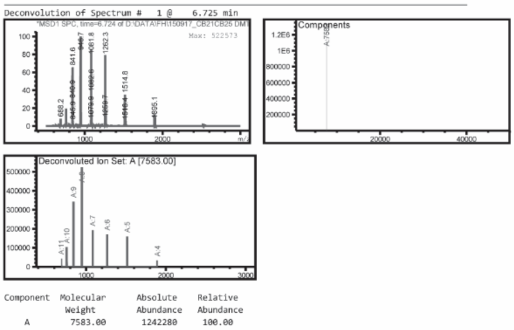

**22** : LNA/DNA mixmer of sequence -45-63. Synthesized at Exiqon (now Qiagen).

| | Name | Seq. 5'-3' | Target | Chemistry | MW found (g/mol) | MW calc. (g/mol) | Δm (%) | UV purity (%) |
|---|---|---|---|---|---|---|---|---|
| **ORN1** | **23** | ag$^{m}$cag$^{m}$c$^{m}$ctgagaaatgtttt$^{m}$ct | -44-65 | Full MOE-PS | 8772.96 | 8770.6 | 0.03 | 98.4 |

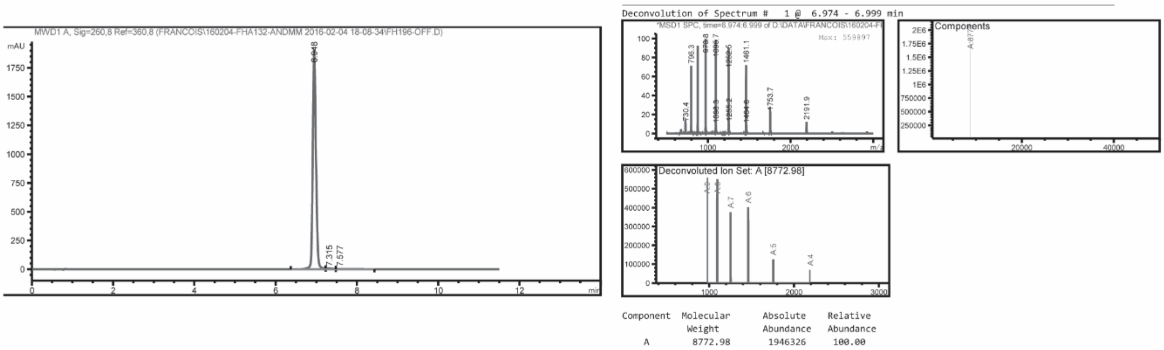

| | Name | Seq. 5'-3' | Target | Chemistry | MW found (g/mol) | MW calc. (g/mol) | Δm (%) | UV purity (%) |
|---|---|---|---|---|---|---|---|---|
| **ORN2** | **24** | $^{m}$cttag$^{m}$cag$^{m}$c$^{m}$ctgagaaatgttt | -41-62 | Full MOE-PS | 8771.37 | 8770.6 | 0.01 | 98.6 |

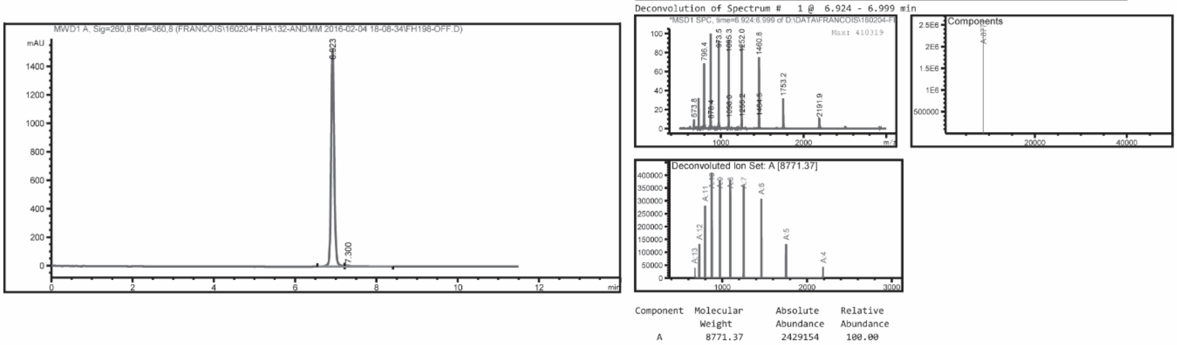

| | Name | Seq. 5'-3' | Target | Chemistry | MW found (g/mol) | MW calc. (g/mol) | Δm (%) | UV purity (%) |
|---|---|---|---|---|---|---|---|---|
| **ORN3** | **25** | $^{m}$cttag$^{m}$cag$^{m}$c$^{m}$ctgagaaatgtttt$^{m}$ct | -41-65 | Full MOE-PS | 9953.08 | 9952.4 | 0.01 | 99.6 |

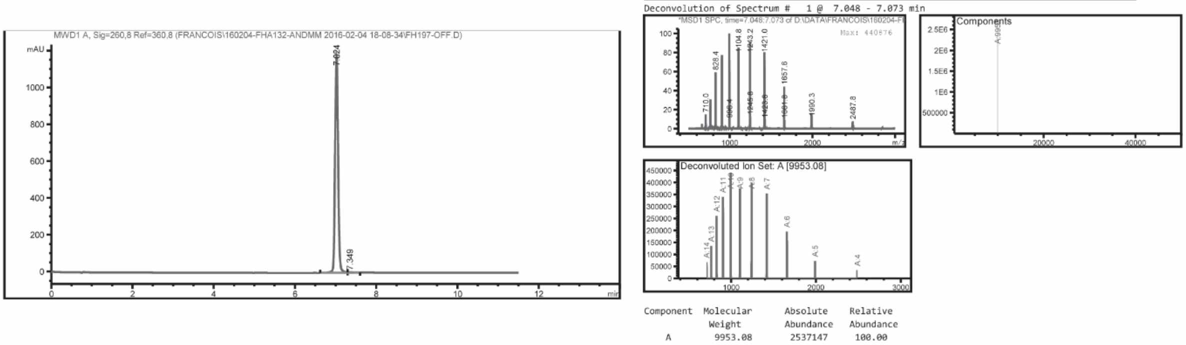

**Neg Ctrl 2**

| Name | Seq. 5'-3' | Target | Chemistry | MW found (g/mol) | MW calc. (g/mol) | Δm (%) | UV purity (%) |
|---|---|---|---|---|---|---|---|
| **26** | gt$^m$cgttagaa$^m$ctta$^m$ctgga$^m$cta | none | Full MOE-PS | 8771.42 | 8770.6 | 0.01 | >99 |

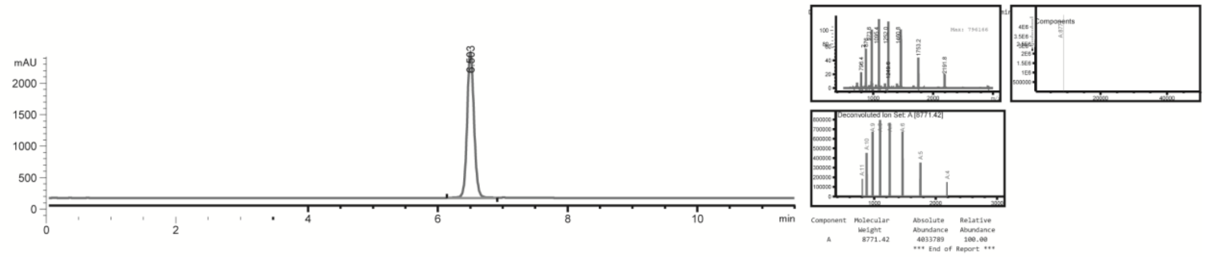

**Neg Ctrl 3**

| Name | Seq. 5'-3' | Target | Chemistry | MW found (g/mol) | MW calc. (g/mol) | Δm (%) | UV purity (%) |
|---|---|---|---|---|---|---|---|
| **27** | ata$^m$cg$^m$cgtattata$^m$cg$^m$cgattaa$^m$cga$^m$c | none | Full MOE-PS | 10768.1 | 10767 | 0.01 | 94 |

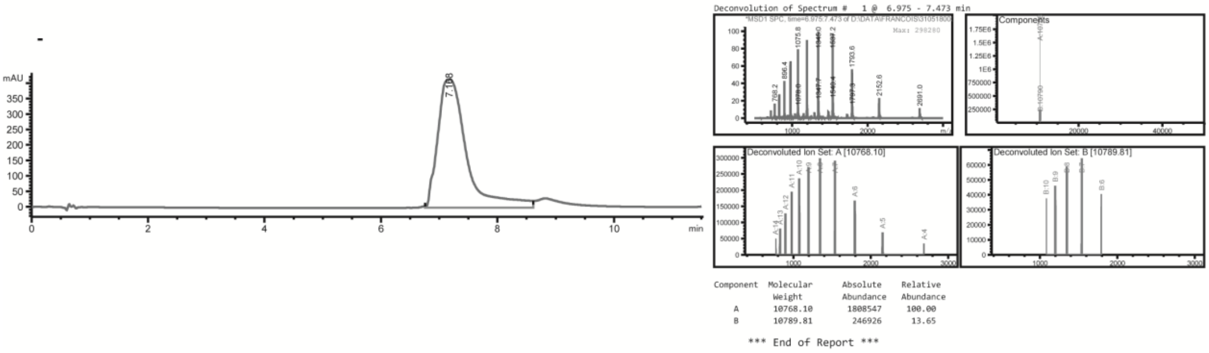

**Figure S2. Oligonucleotide analytics of CL-qPCR primers**

Samples were analysed by LC-MS using conditions described above. Cy3 azide and BHQ azide moieties were coupled on oligonucleotide sequences with 2'-*O*-propargyl modifications according to an in-house protocol (1).

| Name | Seq. 5'-3' | Chemistry | MW found (g/mol) | MW calc. (g/mol) | Δm (%) | UV purity (%) |
|---|---|---|---|---|---|---|
| **28** | TTAAACCAAGAAAACATTT$_{PS}$ | DNA-PO + terminal PS | 5883.13 | 5883.94 | 0.01 | 97.5 |

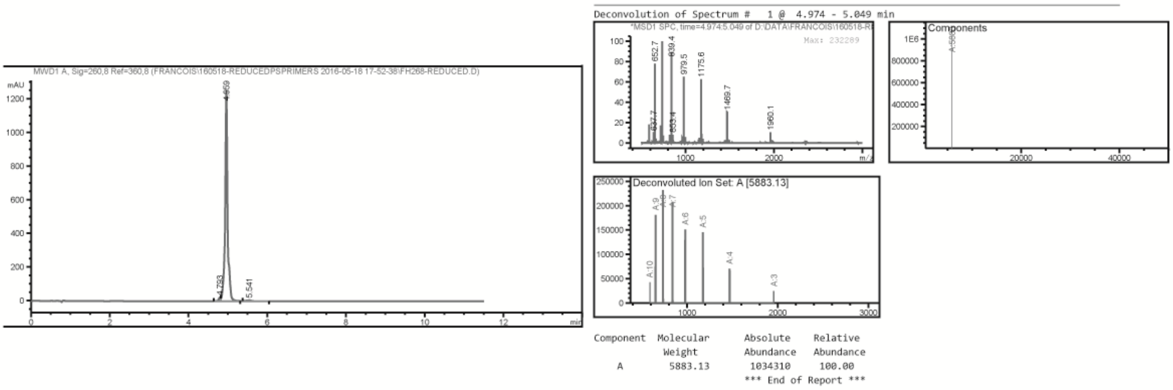

| Name | Seq. 5'-3' | Chemistry | MW found (g/mol) | MW calc. (g/mol) | Δm (%) | UV purity (%) |
|---|---|---|---|---|---|---|
| **29** | ACGTGGTTAGCAGCCTGAGAA | DNA-PO | 6494.9, Na salt | 6495.3 | 0.01 | >99 |

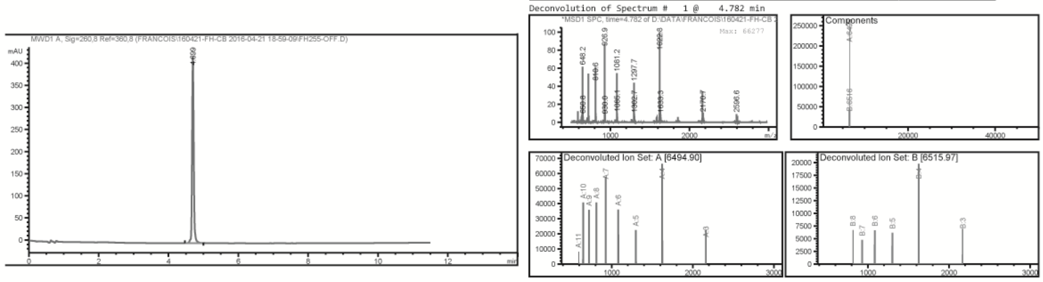

| Component | Molecular Weight | Absolute Abundance | Relative Abundance |
|---|---|---|---|
| A | 6494.90 | 246916 | 100.00 |
| B | 6515.97 | 44969 | 18.21 |

| Name | Seq. 5'-3' | Chemistry | MW found (g/mol) | MW calc. (g/mol) | Δm (%) | UV purity (%) |
|---|---|---|---|---|---|---|
| **30** | $\mathrm{C}_{2'\text{-O-Cy3}}$TCCCTCCCTCGATTTAAACCAAGAAAACAT | DNA-PO Cy3 label | 9967.31 | 9966.65 | 0.01 | >99 |

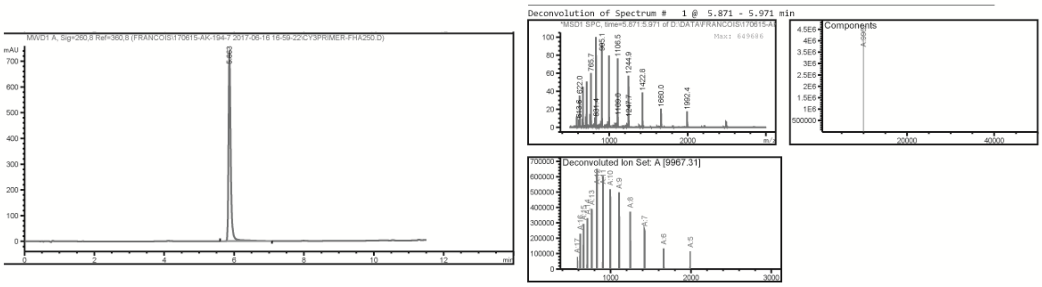

| Component | Molecular Weight | Absolute Abundance | Relative Abundance |
|---|---|---|---|

| Name | Seq. 5'-3' | Chemistry | MW found (g/mol) | MW calc. (g/mol) | Δm (%) | UV purity (%) |
|---|---|---|---|---|---|---|
| **31** | AAATCGAGGGAGGGA$_{2'\text{-O-BHQ}}$G | DNA-PO BHQ label | 5599.23 | 5598.15 | 0.02 | >99 |

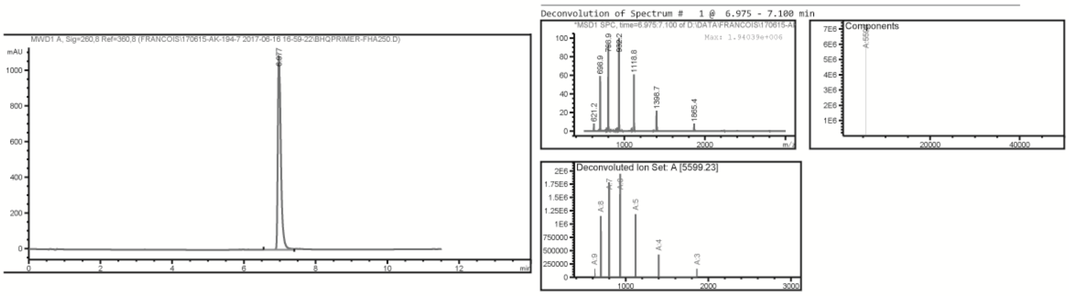

| Component | Molecular Weight | Absolute Abundance | Relative Abundance |
|---|---|---|---|
| A | 5599.23 | 6753415 | 100.00 |

| Name | Seq. 5'-3' | Chemistry | MW found (g/mol) | MW calc. (g/mol) | Δm (%) | UV purity (%) |
|---|---|---|---|---|---|---|
| **32** | BPS-CTCAGGCTGCTAACCACGT | DNA-PO 5' BPS label | 5964.16 | 5966.06 | 0.03 | >99 |

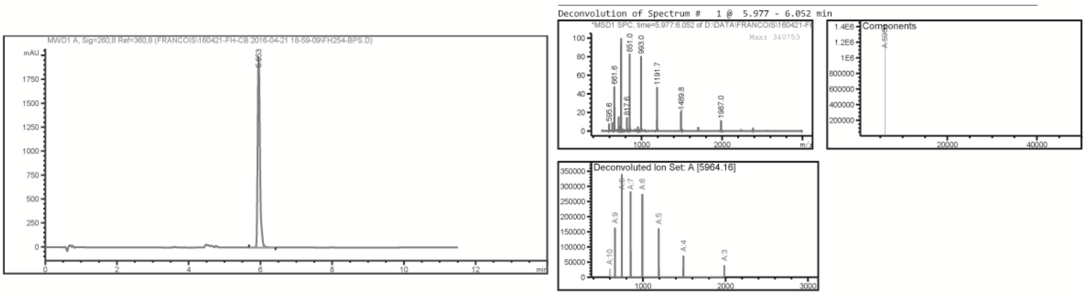

| Component | Molecular Weight | Absolute Abundance | Relative Abundance |
|---|---|---|---|
| A | 5964.16 | 1321300 | 100.00 |

**Figure S3. Analytics of in-vivo batches of ORN1, ORN$^{st}$, ORN$^{84}$ and ORN$^{ch}$**

Oligonucleotide solutions were made in 0.3 M sodium acetate (ThermoFisher #AM9740) after RP-HPLC purification. Two rounds of ethanol precipitation were conducted to generate the compound as a sodium salt. Final dissolution was made in 0.9 % saline (Sigma-Aldrich #S8776), directly followed by a sterile filtration through 0.2 µm filter. An aliquot was sampled under sterile conditions to determine product concentration by $A_{260}$ measurement and purity by LC-MS (Agilent 1200/6130 system) equipped with a Waters Acquity OST C-18 column.

**ORN1**

| | |
|---|---|
| MW calc. (g/mol) | 8770.60 |
| MW found (g/mol) | 8772.72 |
| Δm (%) | 0.02 |
| UV purity (%) | >99 |

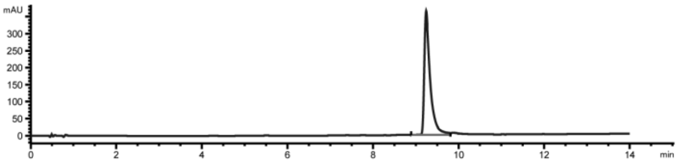

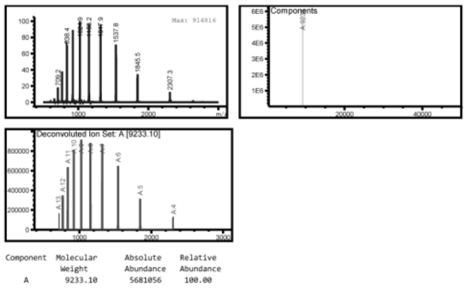

**ORN$^{84}$**

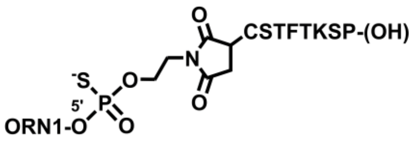

| | |
|---|---|
| MW calc. (g/mol) | 9859.15 |
| MW found (g/mol) | 9860.50 |
| Δm (%) | 0.01 |
| UV purity (%) | >99 |

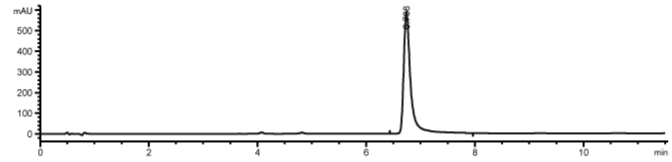

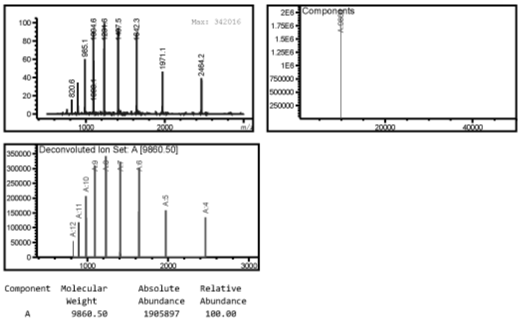

**ORN$^{st}$**

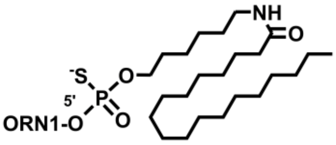

| | |
|---|---|
| MW calc. (g/mol) | 9233.32 |
| MW found (g/mol) | 9233.10 |
| Δm (%) | 0.01 |
| UV purity (%) | >99 |

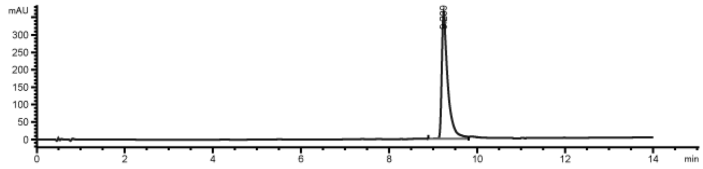

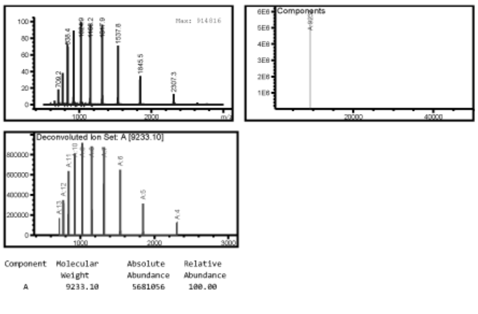

**ORN$^{ch}$**

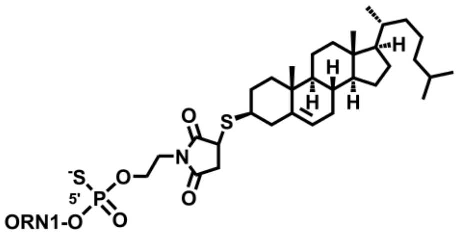

| MW calc. (g/mol) | 9392.42 |
|---|---|
| MW found (g/mol) | 9393.54 + Na ion |
| $\Delta$m (%) | 0.01 |
| UV purity (%) | >99 |

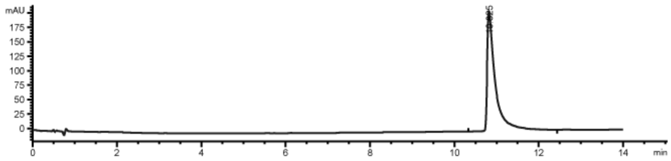

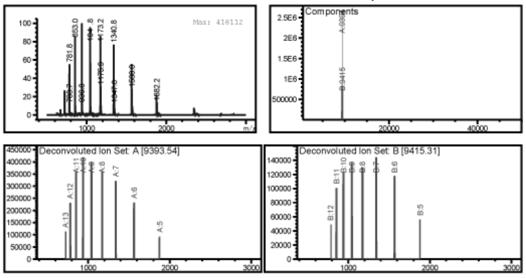

| Component | Molecular Weight | Absolute Abundance | Relative Abundance |
|---|---|---|---|
| A | 9393.54 | 2492164 | 100.00 |
| B | 9415.31 | 834157 | 33.47 |

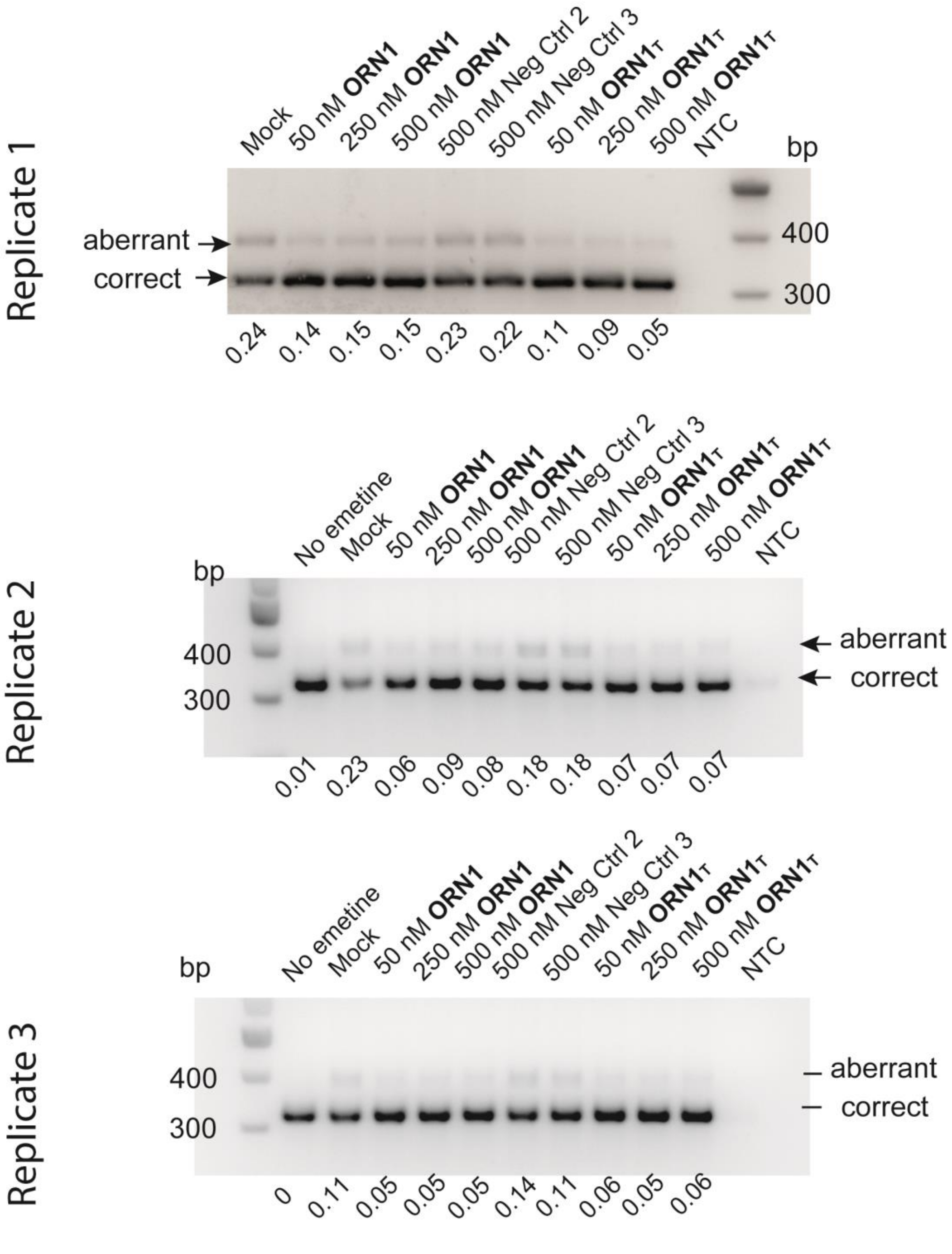

**Figure S4. Gel replicates for *FECH* endogenous splicing correction in K562 cells.**

K562 cells were were plated at 200.000 cells / well in a 24-well plate in 0.5 ml IMDE containing 10 % fetal bovine serum and 4 mM glutamine. Oligonucleotides were transfected using Lipofectamine™ 2000. 24 hours prior to lysis and RNA extraction, emetine was added in each desired well at a final concentration of 3 µM. RNA extraction was carried out using Qiagen RNEasy kit according to manufacturer's instructions. cDNA was generated using PrimeScript RT kit (Takara #RR037A). PCR primers for amplification of human *FECH* transcripts were used. PCR products were loaded on a 2% agarose gel and bands quantified using ImageJ software. Ratios were expressed as: intensity of the aberrant amplicon band / (intensities of aberrant + correct bands).

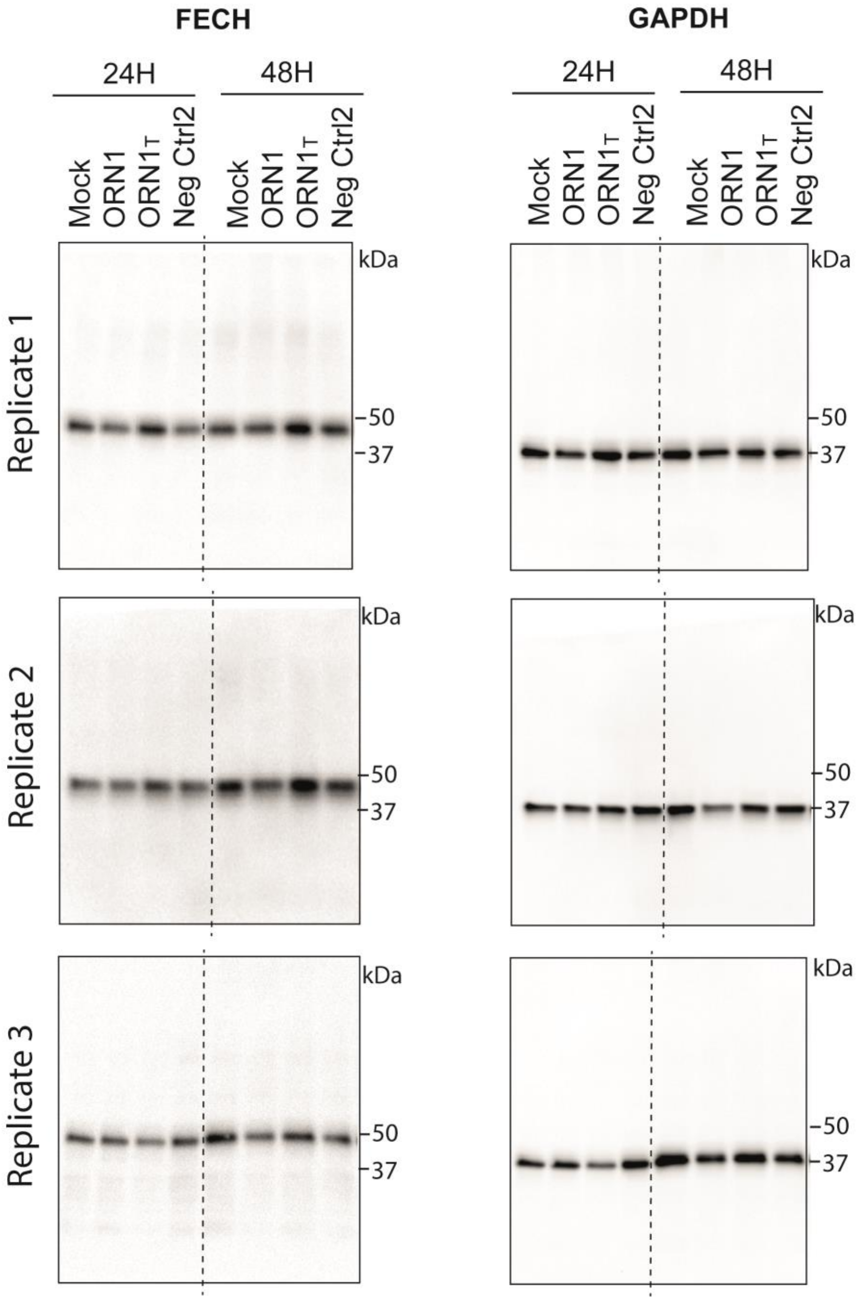

**Figure S5. Western Blot replicates for FECH endogenous splicing correction in K562 cells.**

K562 cells were were plated at 100.000 cells / well in a 24-well plate in 1 ml IMDE containing 10 % fetal bovine serum and 4 mM glutamine. Oligonucleotides were transfected using Lipofectamine™ 2000. Cells were lysed 24 or 48 h after treatment. Protein concentration was assessed with the Pierce BCA protein assay kit (ThermoFisher Scientific). Samples were loaded onto 4–20% pre-cast TGX gels (BioRad) for electrophoresis. The transfer of protein onto PVDF membranes (Roche) was carried out for 90 min at 25 W. Membranes were blocked with 3% BSA and were incubated with the primary antibody overnight at 4 °C. Membranes were washed with 0.05% PBS-T and reacted with the secondary antibody for 1 h in 5% milk. Binding was detected with the ECL Prime reagent (GE Life Sciences) and quantified using the ImageJ software. Graphical output and statistical analysis (Anova) were generated with GraphPad Prism software.

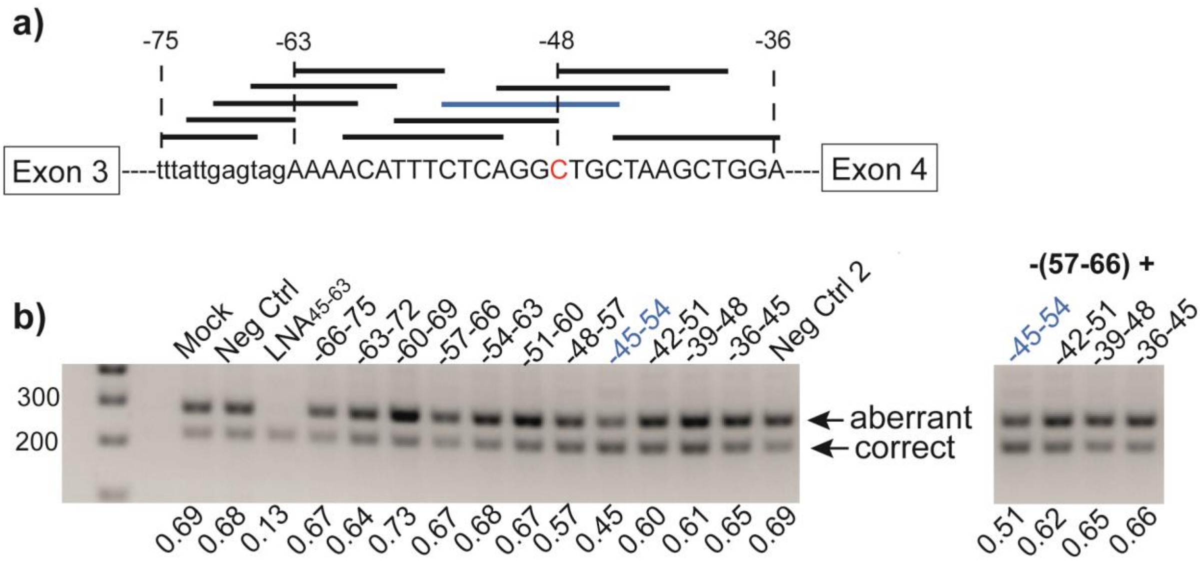

**Figure S6. Binding the c.315-48C SNP but not the aberrant splice site is sufficient for splicing correction.**

**a)** Binding sites for ten 10-nt SSOs in the regions of the -63-splice site and the -48 polymorphism. **b)** Agarose gel showing relative amounts of aberrant and correct transcripts from treatment at 50 nM concentration of COS-7 cells expressing a FECH-C minigene with short SSOs; positive control is -45-63 LNA-SSO. Splicing correction obtained in the minigene experiment with single oligonucleotide treatment (left-hand side) or cotreatments (right-hand side).

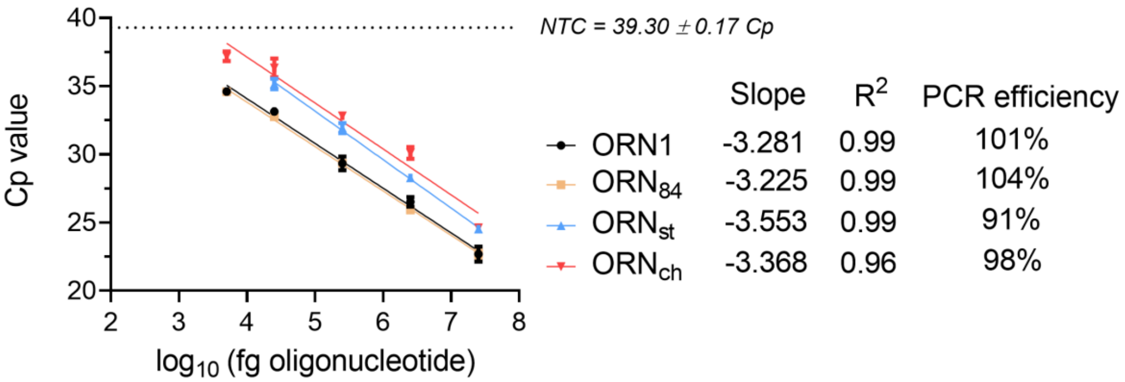

**Figure S7. Calibration curves of oligonucleotide conjugates in kidney lysate**

ORN1 and its conjugated forms were spiked in kidney lysate diluted at 1/750 in nanopure water and serially diluted. Chemical ligation was run for 1 h at 33°C. qPCR was run on a LC480 device (Roche) with technical triplicates for each datapoint. Cp values were calculated by the onboard software. Slopes and $R^2$ values were calculated with GraphPad Prism v7. software. PCR efficiency values were calculated with the web qPCR Efficiency calculator from ThermoFisher Scientific.

#### *In vivo* metabolism study in mouse tissues: mass spectrometry data (Figures S8 to S11)

Grinded tissues were suspended in 10 volumes/weight OTX Lysis Buffer, briefly needle-sonicated and centrifuged for 30 s at 14000 rpm. Supernatants were collected for oligonucleotide clean-up and directly processed with the Clarity® OTX Extraction kit (Phenomenex #KS0-9253), according to manufacturer's procedure for tissues with two modifications: formulation of all buffers with ammonium acetate instead of sodium phosphate; and dissolution of the final eluate in 100 µl of TE buffer pH 8.0 instead of water after evaporation to dryness. Optimization of the protocol with liver samples from mice treated with cholesteryl conjugate **ORN$^{ch}$** comprised a proteinase K pre-treatment of the tissue to ensure recovery of the oligonucleotide (this pre-treatment was used for all **ORN1$^{ch}$** -related treatments). Samples were analysed by LC-MS (Agilent 1200/6130 system) equipped with a Waters Acquity OST C-18 column with a gradient of MeOH in 0.4 mM HFIP, 15 mM trimethylamine; flow rate: 0.3 ml/min. Deconvolution of the mass spectra returned molecular weights from which the structures of metabolites were inferred.

#### Figure S8. MS data of ORN1 in tissues

Kidney: oligonucleotide peak at 5.448 min

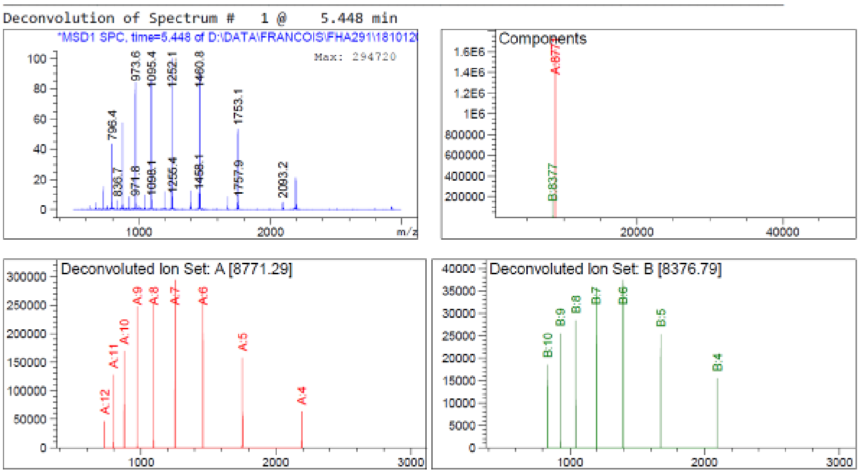

| Component | Molecular Weight | Absolute Abundance | Relative Abundance |
|---|---|---|---|
| A | 8771.29 | 1618850 | 100.00 |
| B | 8376.79 | 181305 | 11.20 |

*** End of Report ***

Liver: Oligonucleotide peak at 5.448 min; n-1 metabolite manually integrated (black arrows)

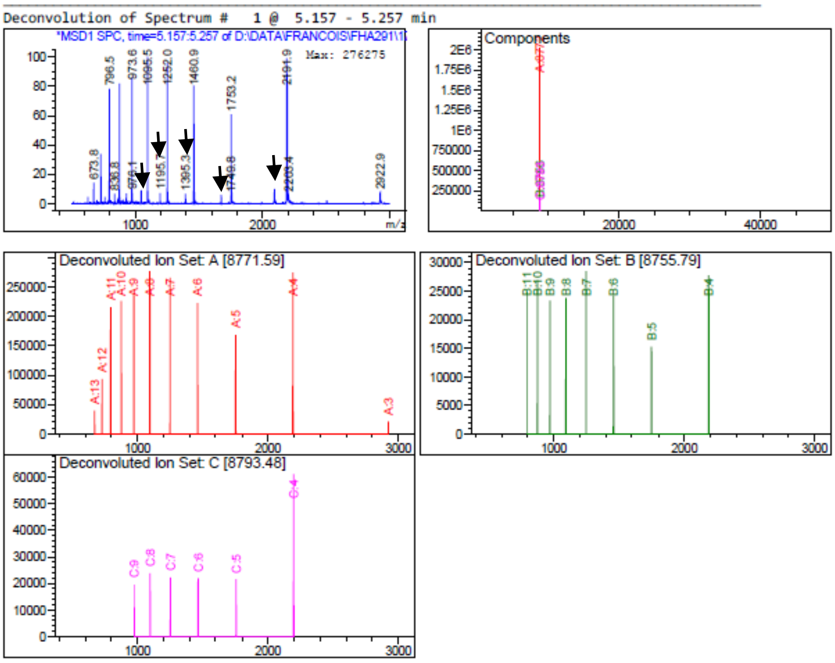

| Component | Molecular Weight | Absolute Abundance | Relative Abundance |
|---|---|---|---|
| A | 8771.59 | 2015827 | 100.00 |
| B | 8755.79 | 189327 | 9.39 |
| C | 8793.48 | 168602 | 8.36 |

*** End of Report ***

Spleen: oligonucleotide peak at 5.556 min; n-1 metabolite manually integrated (black arrows)

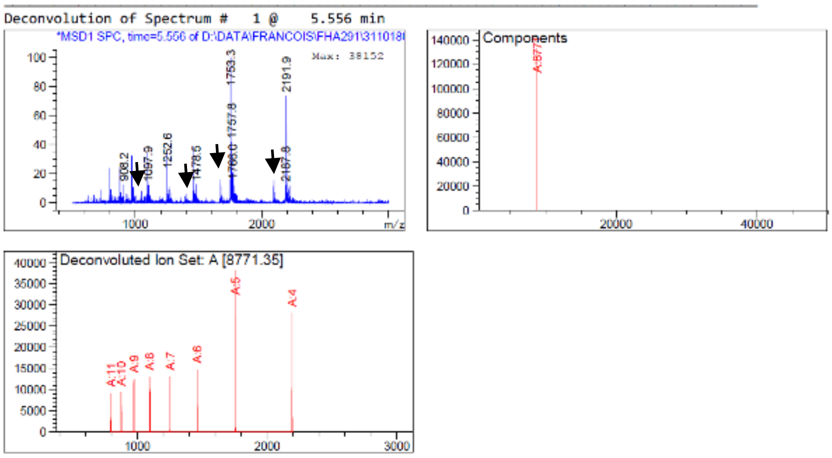

| Component | Molecular Weight | Absolute Abundance | Relative Abundance |
|---|---|---|---|
| A | 8771.35 | 132671 | 100.00 |

*** End of Report ***

Brain: no oligonucleotide detected within sensitivity limits of our equipment

#### Lung: oligonucleotide peak at 5.581 min

Deconvolution of Spectrum # 1 @ 5.581 min

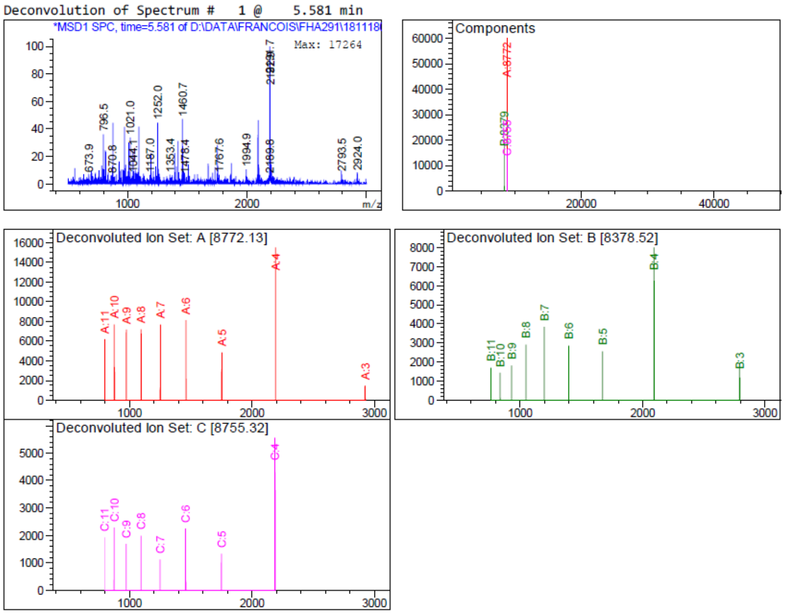

| Component | Molecular Weight | Absolute Abundance | Relative Abundance |
|---|---|---|---|
| A | 8772.13 | 59914 | 100.00 |
| B | 8378.52 | 20764 | 34.66 |
| C | 8755.32 | 16170 | 26.99 |

*** End of Report ***

#### Bone marrow: oligonucleotide peak at 5.581 min

Deconvolution of Spectrum # 1 @ 5.581 min

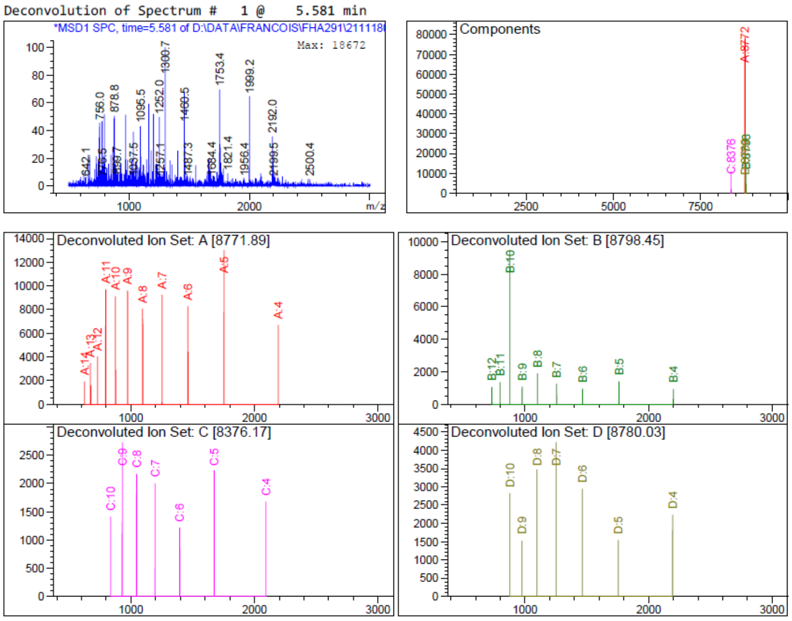

| Component | Molecular Weight | Absolute Abundance | Relative Abundance |
|---|---|---|---|
| A | 8771.89 | 77746 | 100.00 |
| B | 8798.45 | 14106 | 18.14 |
| C | 8376.17 | 11039 | 14.20 |
| D | 8780.03 | 10869 | 13.98 |

*** End of Report ***

#### Figure S9. MS data of ORN$^{st}$ in tissues

Kidney: oligonucleotide peak at 5.456 min

Deconvolution of Spectrum # 1 @ 5.456 min

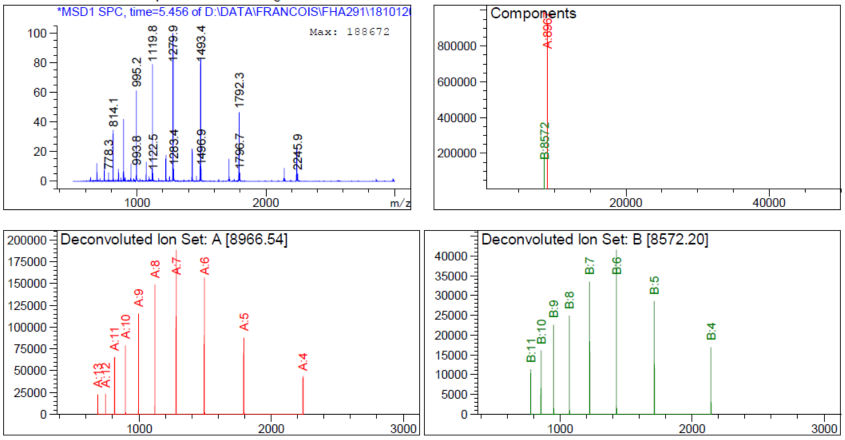

| Component | Molecular Weight | Absolute Abundance | Relative Abundance |
| --- | --- | --- | --- |
| A | 8966.54 | 922098 | 100.00 |
| B | 8572.20 | 189976 | 20.60 |

*** End of Report ***

Liver: two oligonucleotide peaks at 5.381 and 9.292 min

Peak at 5.381 min

Deconvolution of Spectrum # 1 @ 5.381 - 5.431 min

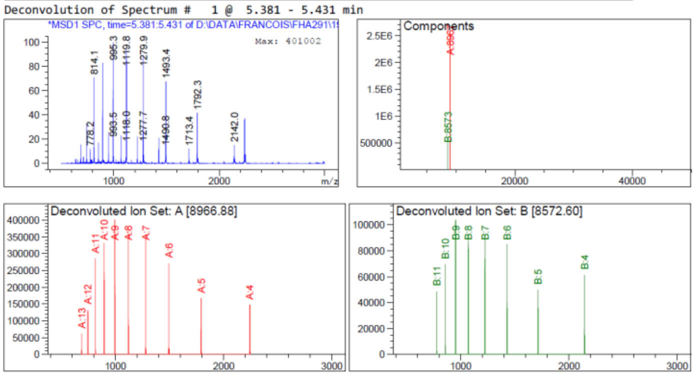

| Component | Molecular Weight | Absolute Abundance | Relative Abundance |
| --- | --- | --- | --- |
| A | 8966.88 | 2509402 | 100.00 |
| B | 8572.60 | 586390 | 23.37 |

Peak at 9.292 min (next page)

Deconvolution of Spectrum # 1 @ 9.292 min

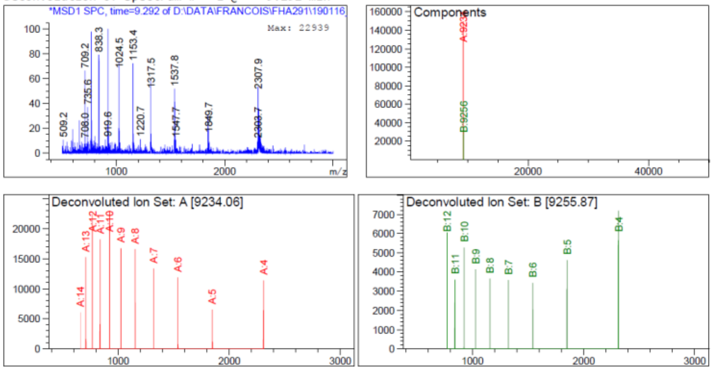

| Component | Molecular Weight | Absolute Abundance | Relative Abundance |
|---|---|---|---|
| A | 9234.06 | 149277 | 100.00 |
| B | 9255.87 | 34066 | 22.82 |

Lung: oligonucleotide peak at 5.506 min

Deconvolution of Spectrum # 1 @ 5.506 min

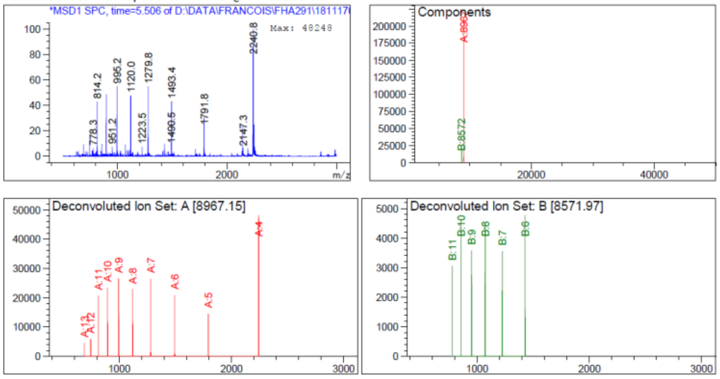

| Component | Molecular Weight | Absolute Abundance | Relative Abundance |
|---|---|---|---|
| A | 8967.15 | 206787 | 100.00 |
| B | 8571.97 | 20236 | 9.79 |

*** End of Report ***

Brain: oligonucleotide peak at 5.481 min

Deconvolution of Spectrum # 1 @ 5.481 - 5.680 min

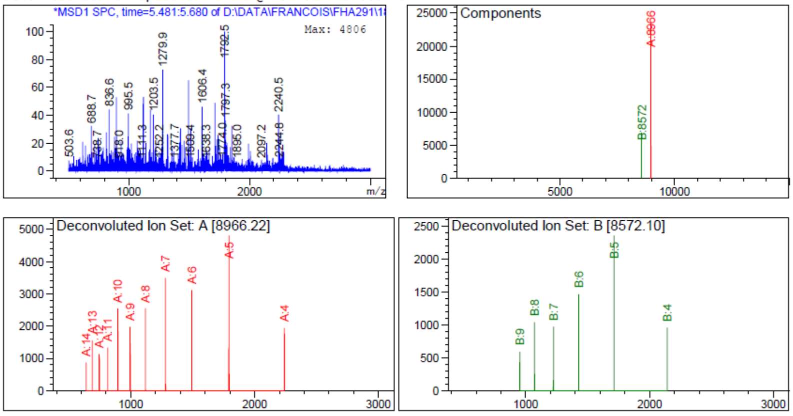

| Component | Molecular Weight | Absolute Abundance | Relative Abundance |
|---|---|---|---|
| A | 8966.22 | 23466 | 100.00 |
| B | 8572.10 | 6763 | 28.82 |

Bone marrow: oligonucleotide peak at 5.597 min

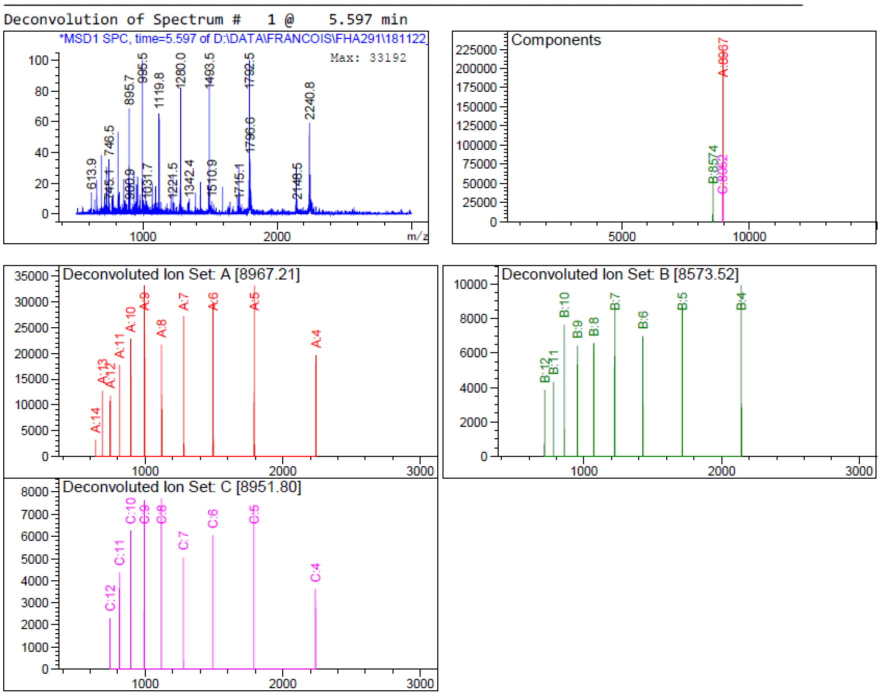

| Component | Molecular Weight | Absolute Abundance | Relative Abundance |
|---|---|---|---|
| A | 8967.21 | 222624 | 100.00 |
| B | 8573.52 | 57352 | 25.76 |
| C | 8951.80 | 42249 | 18.98 |

**Figure S10. MS data of ORN[84] in tissues**

Kidney: oligonucleotide peak at 5.456 min, n-1 metabolite manually integrated (black arrows)

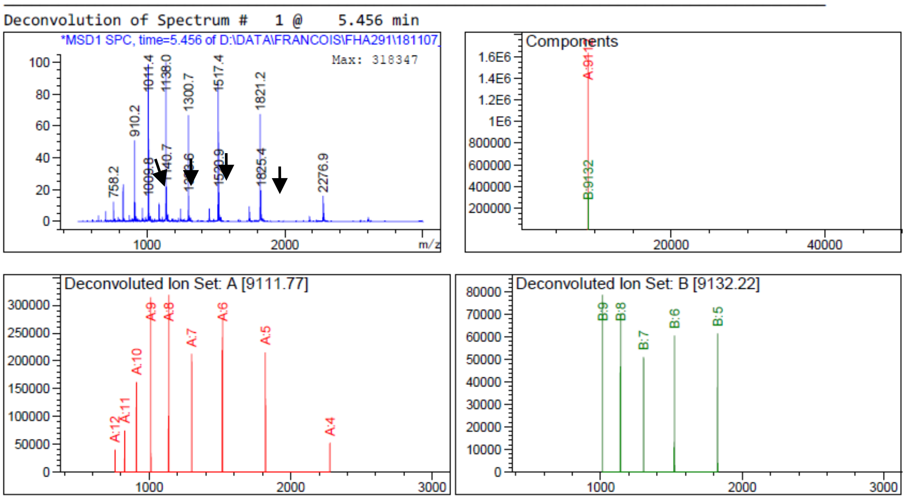

| Component | Molecular Weight | Absolute Abundance | Relative Abundance |
|---|---|---|---|
| A | 9111.77 | 1632938 | 100.00 |
| B | 9132.22 | 314986 | 19.29 |

*** End of Report ***

Liver: oligonucleotide peak at 5.331 min

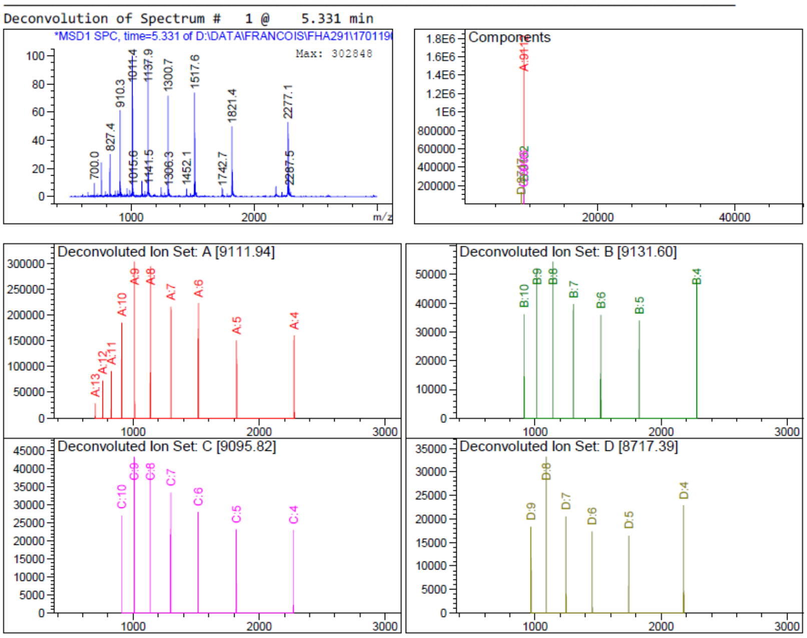

| Component | Molecular Weight | Absolute Abundance | Relative Abundance |
|---|---|---|---|
| A | 9111.94 | 1703340 | 100.00 |
| B | 9131.60 | 285055 | 16.74 |
| C | 9095.82 | 207578 | 12.19 |
| D | 8717.39 | 120787 | 7.09 |

*** End of Report ***

Lung: oligonucleotide peak at 5.431 min

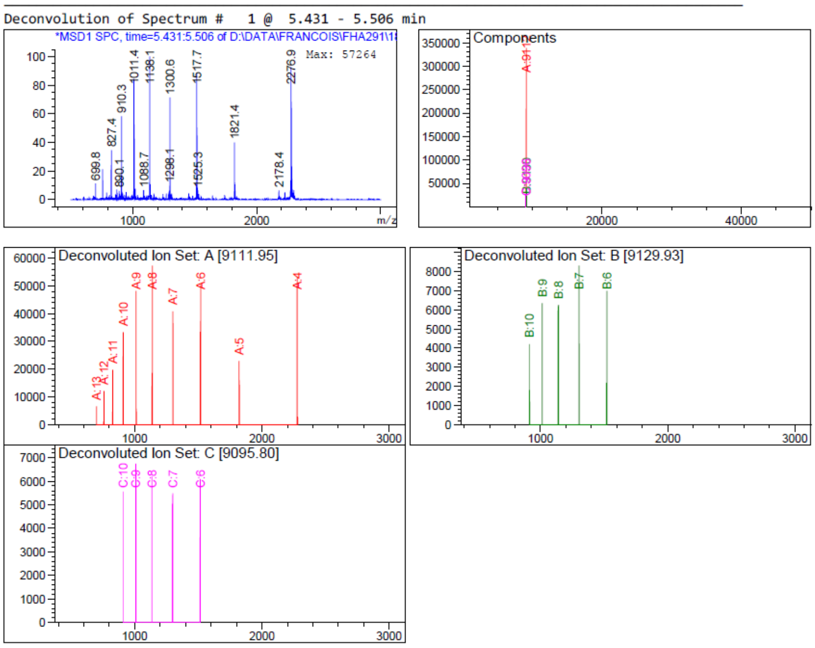

| Component | Molecular Weight | Absolute Abundance | Relative Abundance |
|---|---|---|---|
| A | 9111.95 | 338329 | 100.00 |
| B | 9129.93 | 31450 | 9.30 |
| C | 9095.80 | 27493 | 8.13 |

*** End of Report ***

Bone marrow: oligonucleotide peak at 5.506 min

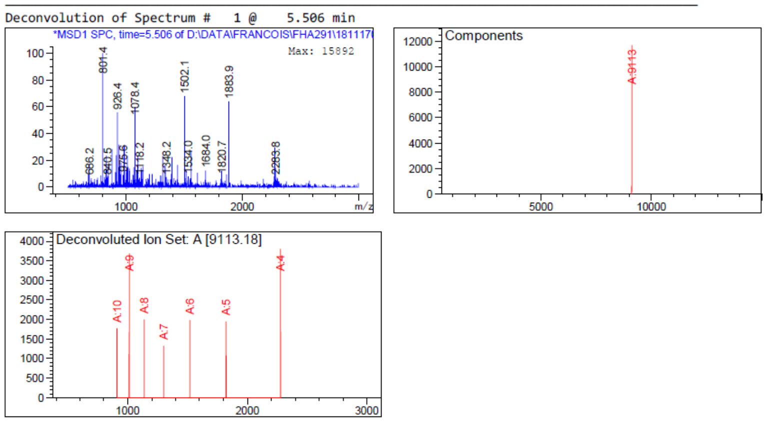

| Component | Molecular Weight | Absolute Abundance | Relative Abundance |
|---|---|---|---|
| A | 9113.18 | 11703 | 100.00 |

Brain: no oligonucleotide detected within sensitivity limits of our equipment

Spleen: oligonucleotide peak at 5.431 min

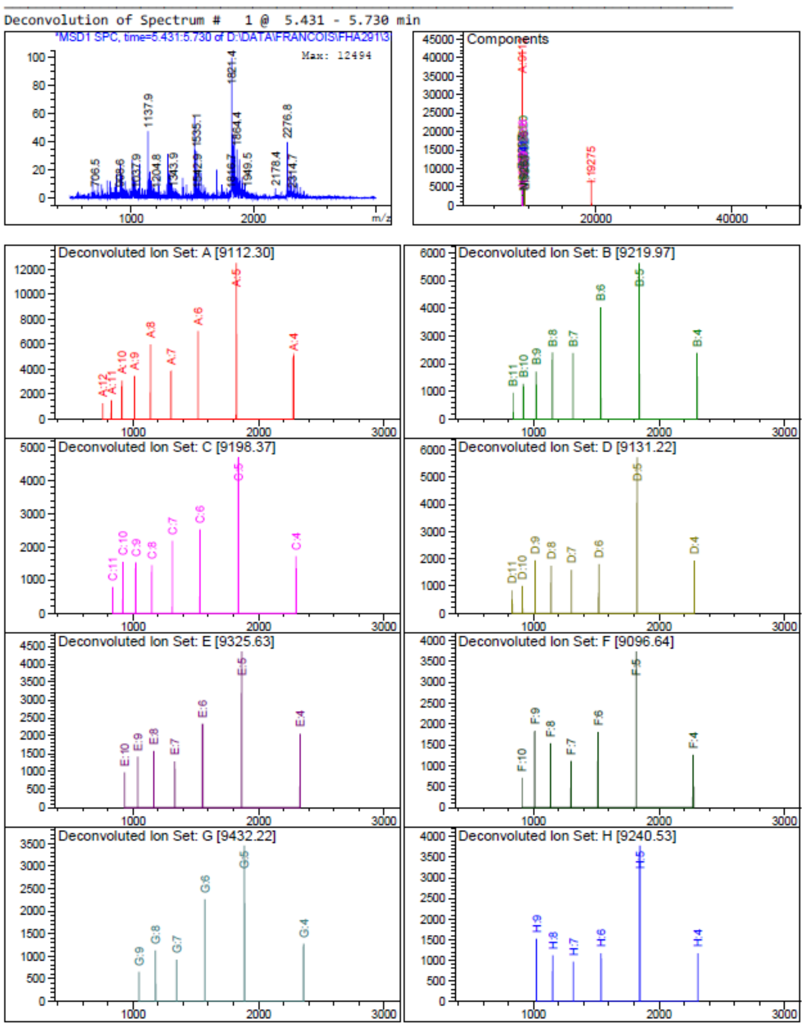

| Component | Molecular Weight | Absolute Abundance | Relative Abundance |
|---|---|---|---|
| A | 9112.30 | 42205 | 100.00 |
| B | 9219.97 | 17292 | 40.97 |
| C | 9198.37 | 16014 | 37.94 |
| D | 9131.22 | 14088 | 33.38 |
| E | 9325.63 | 12336 | 29.23 |
| F | 9096.64 | 11456 | 27.14 |
| G | 9432.22 | 9391 | 22.25 |
| H | 9240.53 | 9088 | 21.53 |

#### Figure S11. MS data of ORN$^{ch}$ in tissues

Kidney: two oligonucleotide peaks at 5.356 min (two main metabolites analyzed) and 11.334 min

Peak at 5.356 min

Deconvolution of Spectrum # 1 @ 5.356 - 5.730 min

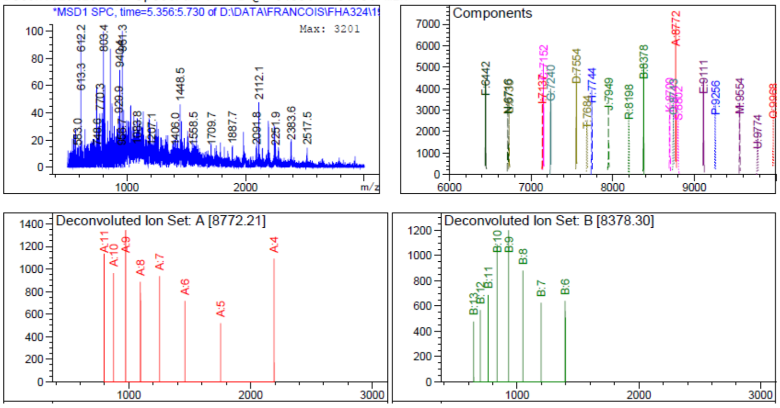

| Component | Molecular Weight | Absolute Abundance | Relative Abundance |
|---|---|---|---|
| A | 8772.21 | 7060 | 100.00 |
| B | 8378.30 | 5210 | 73.80 |

Peak at 11.34 min

Deconvolution of Spectrum # 1 @ 11.334 min

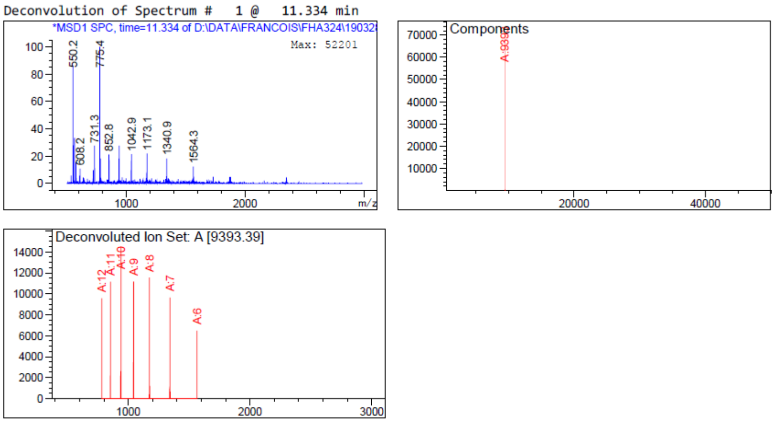

| Component | Molecular Weight | Absolute Abundance | Relative Abundance |
|---|---|---|---|
| A | 9393.39 | 68361 | 100.00 |

*** End of Report ***

Liver: two oligonucleotide peaks at 5.406 min (two main metabolites analyzed) and 10.886 min

Peak at 5.406 min

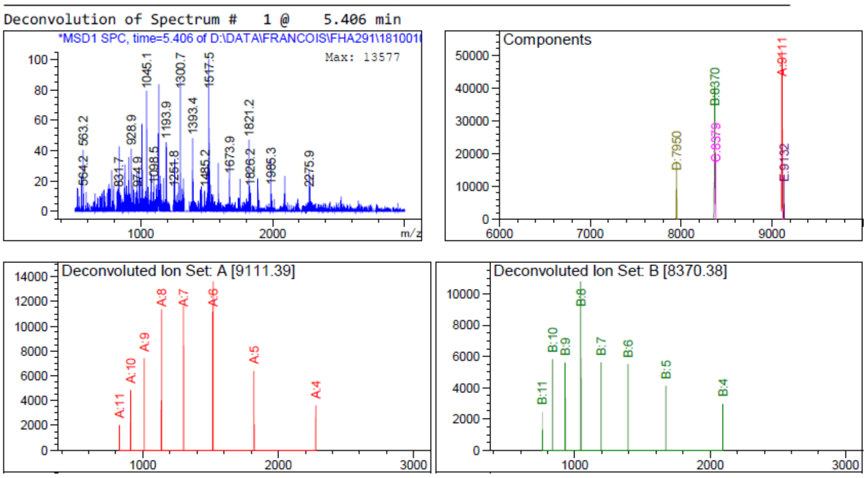

| Component | Molecular Weight | Absolute Abundance | Relative Abundance |
|---|---|---|---|
| A | 9111.39 | 51395 | 100.00 |
| B | 8370.38 | 40597 | 78.99 |

Peak at 10.886 min

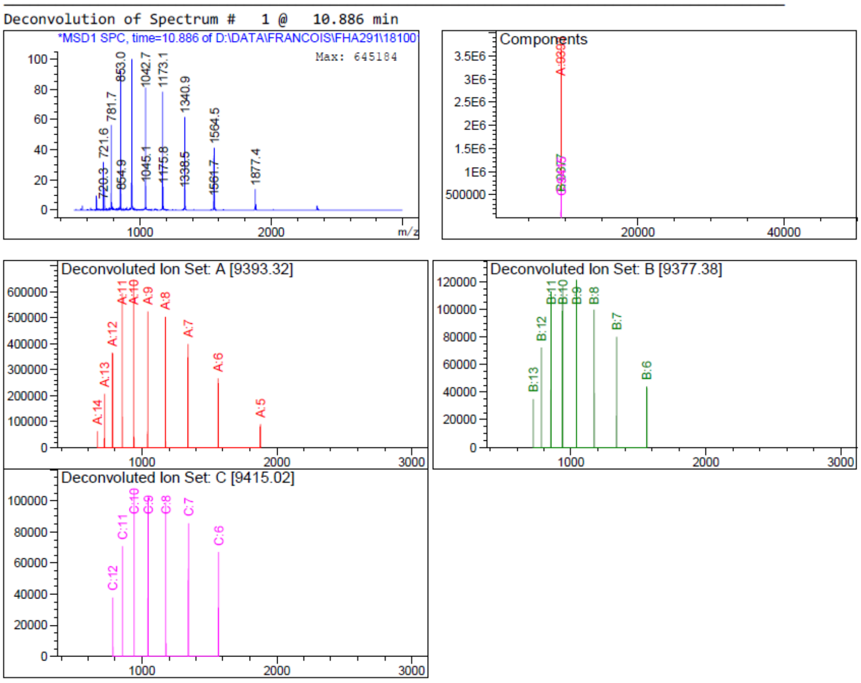

| Component | Molecular Weight | Absolute Abundance | Relative Abundance |
|---|---|---|---|
| A | 9393.32 | 3625731 | 100.00 |
| B | 9377.38 | 662765 | 18.28 |
| C | 9415.02 | 566194 | 15.62 |

*** End of Report ***

Spleen: oligonucleotide peak at 11.409 min

Deconvolution of Spectrum # 1 @ 11.409 - 11.833 min

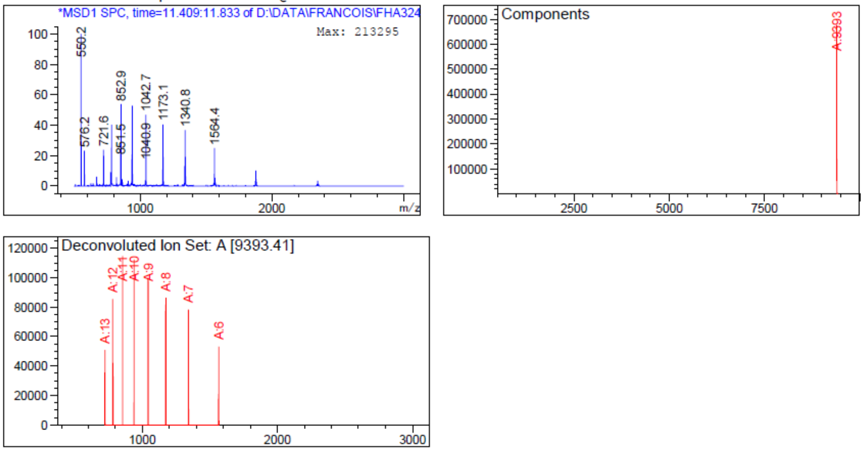

| Component | Molecular Weight | Absolute Abundance | Relative Abundance |
|---|---|---|---|
| A | 9393.41 | 676649 | 100.00 |

*** End of Report ***

Lung: oligonucleotide peak at 11.882 min

Deconvolution of Spectrum # 1 @ 11.882 min

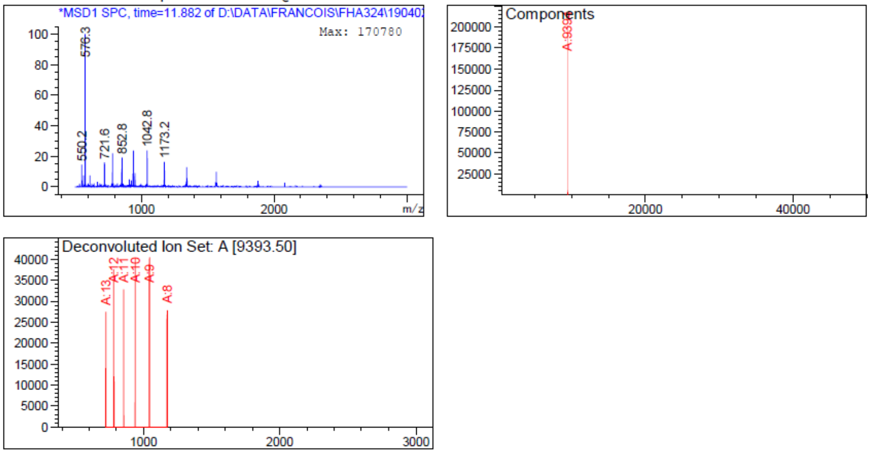

| Component | Molecular Weight | Absolute Abundance | Relative Abundance |
|---|---|---|---|
| A | 9393.50 | 202387 | 100.00 |

*** End of Report ***

Brain: no oligonucleotide detected within sensitivity limits of our equipment

Bone marrow: two oligonucleotide peaks at 5.507 min and 11.884 min (next page)

Peak at 5.507 min

Deconvolution of Spectrum # 1 @ 5.507 min

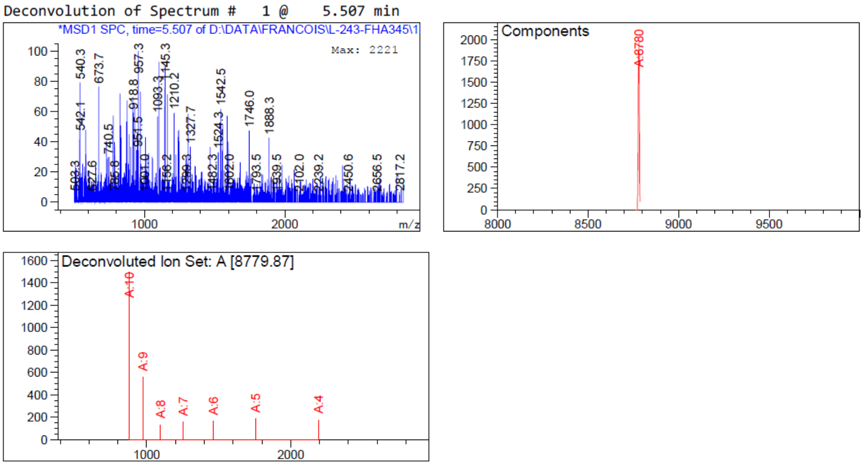

| Component | Molecular Weight | Absolute Abundance | Relative Abundance |
|---|---|---|---|
| A | 8779.87 | 1966 | 100.00 |

*** End of Report ***

Peak at 11.884 min

Deconvolution of Spectrum # 1 @ 11.884 min

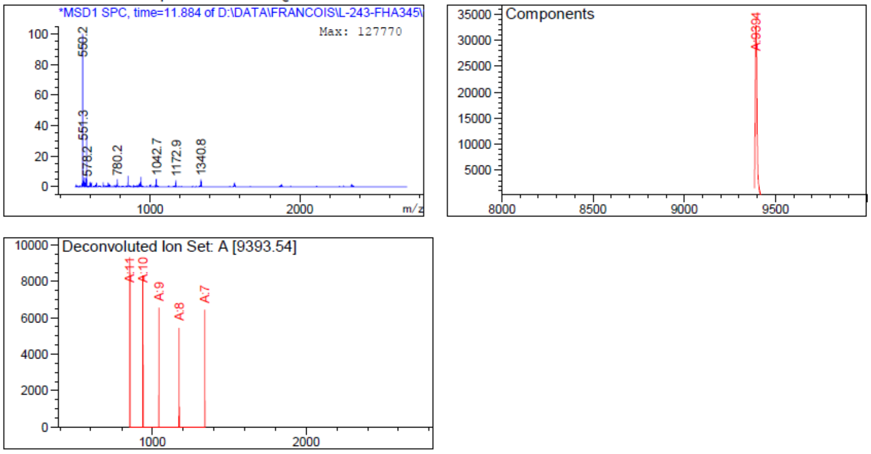

| Component | Molecular Weight | Absolute Abundance | Relative Abundance |
|---|---|---|---|
| A | 9393.54 | 32952 | 100.00 |

*** End of Report ***

#### Supplementary tables

##### Table S1. Library of 10-nt ORNs for minigene screening

Oligonucleotides were produced and characterized as described earlier. Sequences are fully modified MOE-PS chemistry ($^{m}$C refers to 5-methyl cytidine).

| Name | Seq. 5'-3' | Target | MW calc. (g/mol) | MW found (g/mol) | Δm (%) | UV purity (%) |
|---|---|---|---|---|---|---|
| **33** | A$^{m}$CT$^{m}$CAATAAA | -75-66 | 3916.6 | 3917.55 | 0.02 | 93 |
| **34** | T$^{m}$CTA$^{m}$CT$^{m}$CAAT | -72-63 | 3888.5 | 3889.15 | 0.02 | 99 |
| **35** | TTTT$^{m}$CTA$^{m}$CT$^{m}$C | -69-60 | 3870.4 | 3871.55 | 0.03 | 98 |
| **36** | ATGTTTT$^{m}$CTA | -66-57 | 3906.5 | 3905.27 | 0.03 | 97 |
| **37** | GAAATGTTTT | -63-54 | 3941.5 | 3940.80 | 0.02 | >99 |
| **38** | TGAGAAATGT | -60-51 | 3975.6 | 3974.59 | 0.03 | 98 |
| **39** | T$^{m}$CTA$^{m}$CT$^{m}$CAAT | -57-48 | 3973.6 | 3975.05 | 0.04 | 97 |
| **40** | G$^{m}$CAG$^{m}$C$^{m}$CTGAG | -54-45 | 3979.6 | 3979.94 | 0.01 | >99 |
| **41** | TTAG$^{m}$CAG$^{m}$C$^{m}$CT | -51-42 | 3929.5 | 3929.22 | 0.01 | >99 |
| **42** | AG$^{m}$CTTAG$^{m}$CAG | -48-39 | 3964.6 | 3965.59 | 0.02 | >99 |
| **43** | T$^{m}$C$^{m}$CAG$^{m}$CTTAG | -45-36 | 3929.5 | 3929.75 | 0.01 | >99 |
| **44 Neg Ctrl 4** | TAT$^{m}$CATT$^{m}$CA$^{m}$C | Scrambled | 3888.5 | 3888.85 | 0.01 | >99 |

### Supplementary methods

##### Deprotection and purification conditions for lead oligonucleotide ORN1

The CPG was suspended in fresh 25 % ammonia (Sigma Aldrich) and shaken at 45°C for 22 hours. The CPG was filtered, washed with 50 % (aq) EtOH and the liquid fractions were evaporated to dryness prior to dissolution in ultrapure water. After brief centrifugation, the crude DMT-ON oligonucleotide was purified by RP-HPLC (Agilent) with a reverse phase column XBridge OST C18 with a gradient of acetonitrile in 0.1 M aqueous triethylammonium acetate buffer, pH 8. Fractions containing the product were evaporated to dryness and redissolved for detritylation in 50 % (aq) acetic acid. Detritylation was allowed to run under moderate shaking for 1 hour at RT prior to evaporation to dryness and a second round of RP-HPLC purification with the same system. Fractions containing pure product were combined and evaporated to dryness prior to redissolution in ultrapure water.

##### Conjugation on support, deprotection and purification of stearic acid conjugate $ORN_{st}$

The amino C6 modifier (Glen Research #10-1906) was prepared at 0.1M; synthesis was conducted with same parameters as above. The 5' MMT group was deprotected by flushing 3 x 10 ml of Deblock solution through the CPG. The CPG was washed with 2 x 10 ml of acetonitrile and allowed to react with a solution of 160 mg stearic NHS ester and 80 µl DIPEA in 8 ml DMF for 30 min. After washing with 2 x 10 ml of acetonitrile, standard deprotection and purification protocols were applied as for naked compound **ORN1**, however only one round of RP-HPLC was conducted.

##### Deprotection of maleimide-protected form of ORN1 and retro Diels Alder uncaging for conjugation to thiol ligands

The 5'-Maleimide modifier (Glen Research #10-1938) was prepared at 0.1M; synthesis was conducted with same parameters as above. The use of the 5'-maleimide modifier was so far successful with Pac-DNA amidites (2,3) and protocol fine-tuning was required to achieve in the case of MOE oligonucleotides in one hand full deprotection of the nucleobases and cleavage from support, while keeping in the other integrity of the 5' protected maleimide modification. With final conditions, the CPG was suspended in fresh 25 % ammonia (Sigma Aldrich) and shaken at 35°C for 18 hours. For the same reaction time, t temperatures ≥ 40°C led to unacceptable maleimide degradation; temperatures ≤ 30°C to uncomplete nucleobase deprotection. Purification protocol was done as described for **ORN1** but only with one round of RP-HPLC. The 5'-protected maleimide oligonucleotide was analyzed by LC-MS and stored at -20°C at this stage in water; only required amounts were engaged whenever needed for uncaging of the 5'-maleimide and conjugation.

Retro-Diels Alder reaction was conducted using a modified protocol from the literature (3): the 5'-maleimide intermediate was prepared at a concentration of 100 µM in water and microwave-irradiated for 90 minutes at 90°C, followed by immediate concentration in a Speedvac until ≈ 90 % volume reduction was obtained. Evaporation to dryness was avoided. LC-MS injection was routinely carried out to control quality of the retro Diels-Alder deprotection prior to conjugation.

##### Thiocholesteryl conjugate preparation ORN$_{ch}$ from the uncaged maleimide intermediate

A solution of thiocholesterol (Sigma-Aldrich #136115) was prepared in THF at 10 mg/ml and was added to the concentrated solution of maleimide intermediate in water to reach 10 molar eq. of thiocholesterol. The final solvent ratio was adjusted if necessary to get a ratio THF/$H_2O$ of 3:1. The mixture was degassed by argon bubbling and microwave-irradiated for 25 min at 30°C. RP-HPLC purification was then carried out as for compound **ORN1**, but with one round of RP-HPLC only.

##### CD84-binding peptide conjugate preparation ORN$_{84}$ from the uncaged maleimide intermediate

A solution of peptide (Genscript) was prepared in water at 10 mg/ml and was added to the concentrated solution of maleimide intermediate in water to reach 7 molar eq. of peptide. The reaction was buffered with TEAA at pH 7 and allowed to run for 30 minutes at RT. Product **ORN$^{84}$** was purified by RP-HPLC on the same system as for **ORN1** but with milder conditions: 4 ml/min, 40°C and a gradient of acetonitrile in 0.1 M TEAA buffer pH 6.8.
